# Sensory conflict disrupts circadian rhythms in the sea anemone *Nematostella vectensis*

**DOI:** 10.1101/2022.04.11.487933

**Authors:** Cory A. Berger, Ann M. Tarrant

## Abstract

Circadian clocks infer time of day by integrating information from cyclic environmental factors called zeitgebers, including light and temperature. Single zeitgebers entrain circadian rhythms, but few studies have addressed how multiple, simultaneous zeitgeber cycles interact to affect clock behavior. Misalignment between zeitgebers (“sensory conflict”) can disrupt circadian rhythms, or alternatively clocks may privilege information from one zeitgeber over another. Existing studies are limited in the range of tested zeitgeber relationships, and also in their taxonomic breadth, which is restricted to insects and vertebrates among animals. Here, we show that temperature cycles entrain circadian locomotor rhythms in *Nematostella vectensis*, a model system for cnidarian circadian biology. We then conduct behavioral experiments across a comprehensive range of light and temperature cycles. *Nematostella*’s circadian behavior is disrupted by chronic sensory conflict, including disruption of the endogenous clock itself rather than a simple masking effect. Sensory conflict also disrupts the rhythmic transcriptome, with numerous genes losing rhythmic expression. However, many metabolic genes remained rhythmic and in-phase with temperature, and other genes even gained rhythmicity, implying that some rhythmic metabolic processes persist even when behavior is disrupted. Our results show that a cnidarian clock relies on information from light and temperature, rather than prioritizing one zeitgeber over the other. Although we identify limits to the clock’s ability to integrate conflicting sensory information, there is also a surprising robustness of behavioral and transcriptional rhythmicity.

## Introduction

Nearly all organisms use circadian clocks to synchronize their behavior, physiology, and metabolism to ∼24h environmental cycles. Factors that synchronize circadian clocks are called zeitgebers, and include light, temperature, and food availability (***Aschoff, 1960***). In nature, clocks must integrate information from many zeitgebers simultaneously to infer coherent clock outputs, although most of our knowledge of circadian clocks comes from single-zeitgeber experiments. During “sensory conflict,” when zeitgebers give conflicting information to the clock, circadian rhythms can either be disrupted (***Harper et al., 2016***) or preferentially follow a single zeitgeber (***Harper et al., 2017***). By defining conditions under which normal clock output is possible or disrupted, studies of sensory conflict help us understand how environmental information is transmitted into rhythmic behavior, which is a central goal of circadian biology (reviewed for insects in ***Rivas et al., 2016; Somers et al., 2018***). Additionally, the behavior of clocks under conflicting zeitgeber regimes is particularly relevant to human society, where factors such as shift work and artificial light regimes are connected to a wide range of metabolic and neurological diseases (***Ehlers et al., 1988; Roenneberg and Merrow, 2016***). However, few studies in any animal have comprehensively characterized clock function in multi-zeitgeber systems, and in general we do not understand how animals behave during sensory conflict.

Current research of sensory conflict is limited by a dearth of comprehensive behavioral experiments. The small literature of sensory conflict studies has examined conflicting light and temperature cycles in insects (***Miyasako et al., 2007; Currie et al., 2009; Watari and Tanaka, 2010; Yoshii et al., 2010; Nikhil et al., 2014; Harper et al., 2016, 2017; Rivas et al., 2018; Kaniewska et al., 2020***), vertebrates (***Firth and Kennaway, 1989; Valenciano et al., 1997; Moyer et al., 1997; Firth et al., 1999***), protists (***Bruce, 1960***), and cyanobacteria (***Lin et al., 1999***). There have also been studies of light and time-restricted feeding in vertebrates (***Javier Sánchez-Vázquez et al., 1995; Challet et al., 1998; Reierth and Stokkan, 1998; Lague and Reebs, 2000***). However, many of these studies did not compare more than two possible phase relationships between zeitgebers, precluding characterization of the potentially non-linear relationship between an entrained rhythm and two entraining cues. It is necessary to test finer degrees of misalignment between zeitgebers to understand the sensitivity of clock outputs to environmental conditions.

Another critical knowledge gap is how sensory conflict affects rhythmic gene expression, which ultimately underlies rhythmic physiology and behavior (***Yeung and Naef, 2018***). Animal clocks function via transcription-translation feedback loops in which transcription factors activate the expression of their own repressors, and also regulate transcription of downstream genes (***Partch et al., 2014; Dubowy and Sehgal, 2017***). Although one study measured the expression of individual clock genes during sensory conflict (***Rivas et al., 2018***), the effects on downstream gene expression are entirely unknown. Animals exposed to arrhythmic conditions, such as mice in feeding experiments (***Greenwell et al., 2019***) or cavefish over evolutionary time (***Mack et al., 2021***), often express fewer rhythmic transcripts. Transcriptome-scale studies are needed to test whether sensory conflict similarly disrupts expression of core clock genes or of downstream clock-controlled genes, resulting in a general loss of rhythmic gene expression.

We address these questions with the model system *Nematostella vectensis*, the starlet sea anemone. Studies of circadian rhythms in Cnidaria, the sister group to Bilateria (***Simion et al., 2017***), are broadly important for understanding how clocks have evolved and diversified in animals. Specifically, cnidarians provide an intriguing system for circadian sensory processing because they lack a central nervous system. In bilaterians, researchers often distinguish central clocks in the brain and peripheral clocks in other tissues, with the central clock acting to synchronize peripheral clocks (***Dibner et al., 2010***). Certain phase relationships between light and temperature cycles disrupt the central clock in *Drosophila* (***Harper et al., 2016***), but not peripheral clocks, which preferentially entrain to light (***Harper et al., 2017***). Thus, sensory conflict might disrupt organism-level behavior because of the differential entrainment of central and peripheral clocks. Cnidarians by definition lack this distinction and therefore provide an important and complementary framework to bilaterian models for studying sensory conflict. *Nematostella* exhibits nocturnal rhythms in locomotion that entrain to artificial light-dark cycles (***Hendricks et al., 2012; Oren et al., 2015***) and to natural environmental conditions (***Tarrant et al., 2019***). Although temperature cycles are a nearly-universal entraining cue for circadian rhythms (e.g ***Somers et al., 1998; Glaser and Stanewsky, 2005; Lahiri et al., 2005; Yoshida et al., 2009; Hart et al., 2021***), no studies have yet investigated thermal entrainment in a non-bilaterian animal.

Here, we show that environmentally realistic temperature cycles synchronize circadian behavior in *Nematostella*, producing activity profiles similar to light-driven rhythms. We then comprehensively test different combinations of light and temperature cycles and show that aligned cycles drive normal rhythmic behavior, while behavior becomes arrhythmic under large degrees of sensory conflict. In particular, we demonstrate the disruption of free-running rhythms, indicating changes to the endogenous clock itself. Using RNA sequencing, we find that sensory conflict substantially alters diel gene expression patterns. This includes both the loss and gain of rhythmic genes, the disruption of co-expressed gene modules related to cellular metabolism, and an overall weakening of correlations between rhythmic genes. However, much of the transcriptome remained rhythmic and in-phase with the temperature cycle. This study extends our understanding of temperature entrainment and integration of multiple zeitgebers to a non-bilaterian model animal, showing that light and temperature interact to synchronize clock outputs, with neither cue totally dominating the other. Thus, normal organismal rhythms are only possible under certain environmental conditions.

## Results

### Temperature cycles drive circadian rhythms in *Nematostella*

We first showed that gradual, environmentally-relevant temperature cycles drive diel locomotor behavior and entrain circadian rhythms in *Nematostella*. Anemones maintained in constant darkness with a temperature cycle that changed gradually from 14–26 °C (1 degree per hour; ***Figure 1***a) exhibited rhythmic diel behavior (n=18, eJTK test, p=3 × 10^−5^; LSP permutation test, *p* < 0.001, ***Figure 2***a). With ZT0 defined as the coldest point of the cycle, peak activity occurred at ZT18. The temperature cycle also entrained free-running rhythms that persisted during constant temperature at 20 °C (LSP, *p* < 0.001, n=23), although the free-running period did not occur at exactly 24 h (eJTK, *p* = 0.99). The proportion of free-running individuals as determined by periodograms is comparable to that of anemones entrained to light cycles in a previous study (35% vs. 33%) (***Tarrant et al., 2019***). 14–26 °C is well within the daily temperature range experienced by the source population of *Nematostella* (***Tarrant et al., 2019; Sachkova et al., 2020***), and we observed similar results for a broader temperature cycle (8–32 °C, ***Figure 2***a) that is also within the range of *in situ* diel variation. We used qRT-PCR to measure the expression of several core circadian genes (*Clock, Cry1a, Cry1b, Cry2*, and *Helt*) over 48 h of the 8–32 °C temperature cycle, and also during free-run. Only *Cry2* was rhythmic during the temperature cycle (LSP, *p* < 0.001), with peak expression at roughly ZT0 (***Figure 2***c). No genes were rhythmic during free-run.

**Figure 1.**
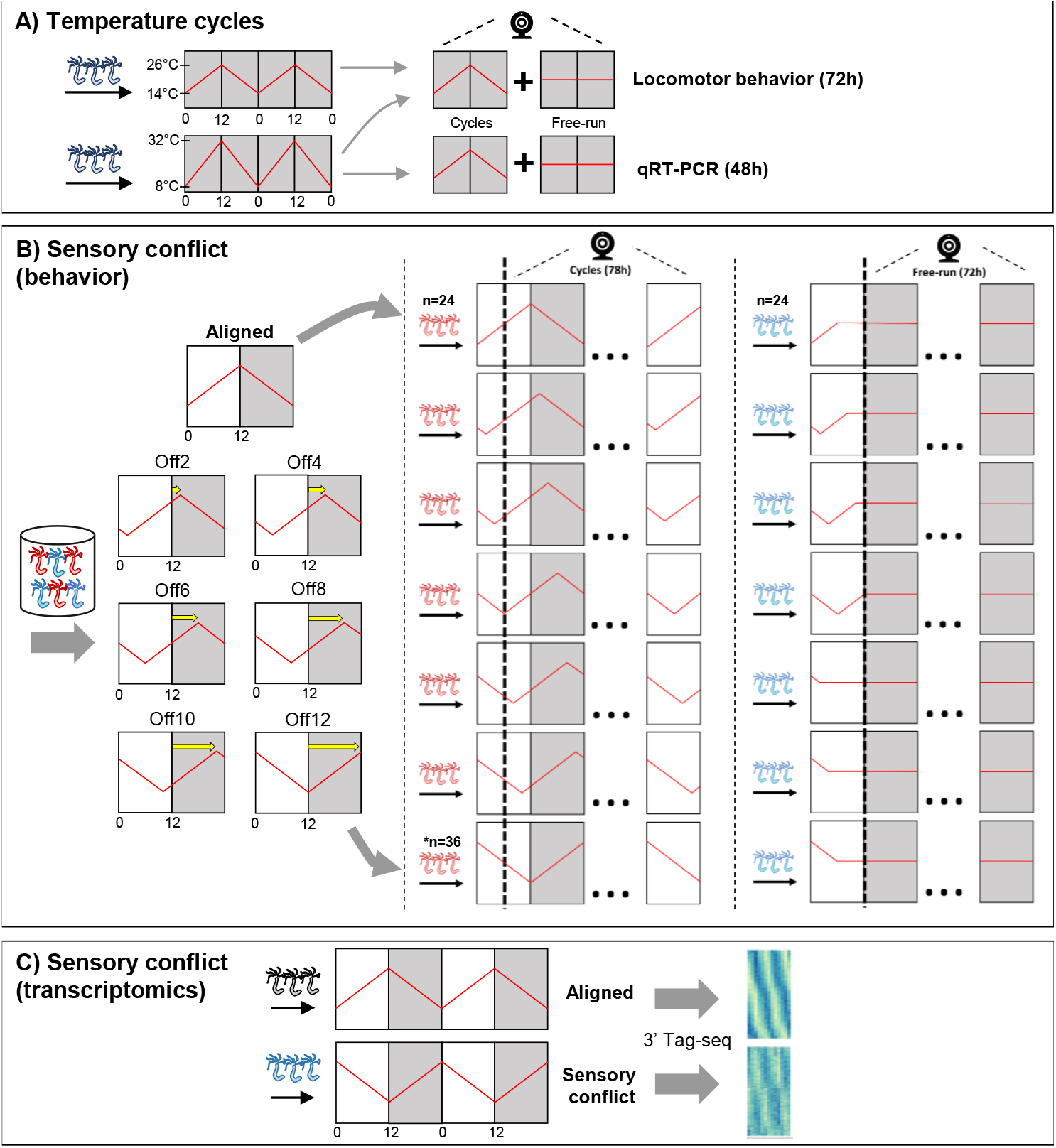
Schematic of the experimental design. **A)** Anemones in constant darkness were entrained to ramped temperature cycles that changed gradually from either 14–26 °C, or from 8–32 °C. ZT0 refers to the coldest time point. Locomotor behavior was recorded during cycling and free-running conditions for both cycles, and qRT-PCR was used to measure the expression of select clock-related genes for the 8–32 °C cycle. **B)** Animals were entrained to 12:12 b light-dark cycles and simultaneous 14–26 °C temperature cycles. In the Aligned (reference) group, the coldest part of the temperature cycle was aligned with lights-on; ZT0 always refers to lights-on. For 6 other groups of anemones, the phase of the temperature cycle was delayed relative to the light cycle in 2h increments up to 12h. Anemones were acclimated to experimental conditions for at least 2 weeks before measuring behavior. For each group, behavior was recorded for 78h during cycles (n=24^a^). Behavior was recorded for separate groups of anemones (n=24) for 72h during free-running (dark, 20 °C). Shaded and unshaded regions represent dark and light periods, respectively. **C)** Gene expression was measured with 3’-anchored Tag-seq during aligned light and temperature cycles, and 12h offset cycles. Anemones were sampled every 4h for 48h (13 time points) simultaneously for both time series. There were n=3 biological replicates at each time point, and each replicate consisted of 5 pooled individuals^b. a^Except 12h offset, which had n=36. ^b^Except one library (at the 9^th^ time point in the Aligned time series), which consisted of 3 pooled individuals

**Figure 2.**
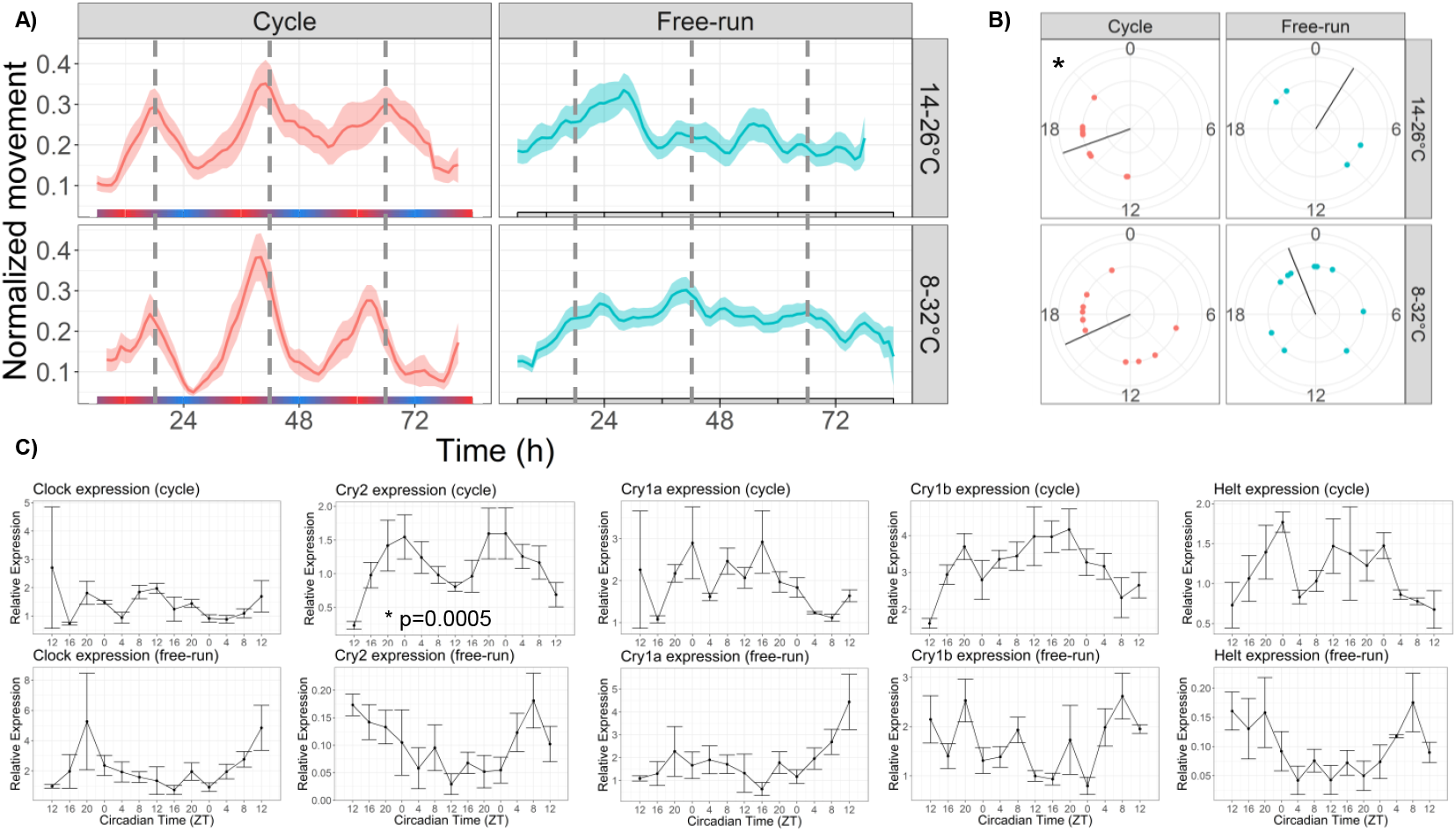
Temperature cycles drive rhythmic locomotor behavior and *Cry2* expression, and weakly entrain circadian behavior. **A)** Mean behavior profiles of anemones in each group over 3 days. Individual locomotor profiles were normalized, averaged, and smoothed (see Methods). Shaded area represents standard error. Left, temperature cycles with scale bar indicating cold (blue) and hot (red) temperatures. Right, free-running at 20 °C. Dashed lines indicate ZT18, the period of peak activity during cycling conditions. **B)** Phases of rhythmic animals calculated by MFourFit. Black line represents circular mean. *:Rayleigh test, p<0.05. **C)** Expression of core circadian genes entrained to a temperature cycle and under free-running conditions. Only *Cry2* was significantly rhythmic (LSP, p<0.05). **Figure 2–source data 1**. Behavioral rhythmicity results.

### Aligned light and temperature cycles entrain robust diel rhythms

We tested how the relationship between light and temperature cycles affects circadian behavior by acclimating groups of anemones to a gradually changing temperature cycle (14–26 °C) simultaneously with a 12:12 light-dark cycle. A schematic of the experimental design is shown in ***Figure 1***b. When light and temperature cycles were aligned such that lights-on corresponded to the coldest part of the temperature cycle (ZT0), *Nematostella* exhibited clear circadian behavior during both cycling (***Figure 3***a; eJTK, p=2 × 10^−5^) and free-running conditions (eJTK, p=7 × 10^−5^). During the cycles, 20/24 individual (83%) had significant 24h rhythms and peak activity occurred at ZT19, consistent with the phases of locomotor activity driven by light (ZT16-20, ***Hendricks et al., 2012; Oren et al., 2015***) and temperature cycles (ZT17-18, ***Figure 2***a). During free-run, 10/24 individuals (42%) had significant 24h rhythms and peak activity occurred at ZT12 (Table 2).

**Figure 3.**
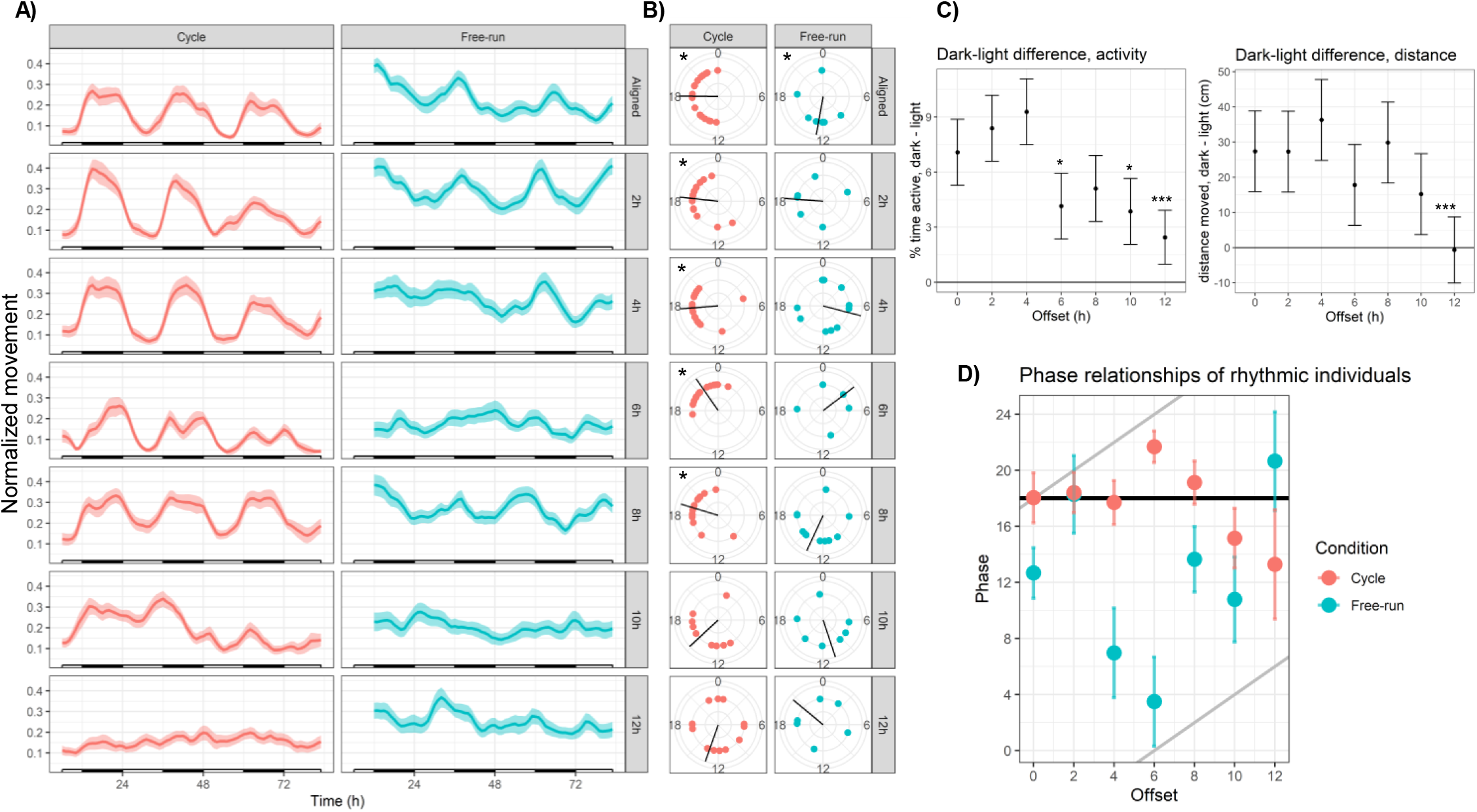
Sensory conflict disrupts circadian locomotor behavior in *Nematostella*. **A)** Mean behavior profiles. Individual locomotor profiles were normalized, averaged, and smoothed (see Methods). Shaded area represents standard error. Left, cycling conditions; white and black bars indicate lights-on and lights-off. Right, free-running at 20 °C and constant darkness; gray and black bars indicate subjective day and night. n=24 for each group except Off12-cycle, where n=36. **B)** Phases of rhythmic animals calculated by MFourFit. Black line represents circular mean. *:Rayleigh test, p<0.05. **C)** Difference, in percentage points, between percent time active per hour during dark and light phases (left), and difference between average distance moved per hour during dark and light phases (right). Asterisks indicate significant difference from Aligned-cycle. *: *p* < 0.05; ***: *p* < 0.001. **D)** Means and standard errors of the phases of rhythmic individuals in each group. Black and grey lines represent expected phases of light-entrained and temperature-entrained rhythms, respectively (see “Relative strengths of light and temperature zeitgebers”).

### Sensory conflict disrupts rhythmic behavior

We tested the effects of long-term entrainment to different zeitgeber regimes by delaying the phase of the temperature cycle relative to the light cycle in 2h increments (***Figure 1***b). Anemones were transferred from aligned zeitgeber cycles to one of 6 offset regimes (2, 4, 6, 8, 10, and 12h offsets) and acclimated for at least two weeks. Within each zeitgeber regime, we recorded the behavior of separate groups of animals during cycling and free-running conditions, resulting in a total of 14 experimental groups including aligned conditions. We use the following notation to refer to experimental groups: Off6-cycle refers to a 6h offset between zeitgebers, recorded during the cycles; Off6-FR refers to behavior recorded during free-run. We quantified the strength of circadian rhythms using several metrics: the maximum power of Lomb-Scargle periodograms; the amplitude of mean time series; the number of individuals with significant 24h rhythms above a strict cutoff (eJTK *p* < 0.001); the relative synchronization (circular variance) of the phases of rhythmic individuals; and the difference in activity between light and dark phases.

Large degrees of sensory conflict (SC) profoundly disrupted rhythmic behavior. During the largest possible misalignment (Off12-Cycle) the normal activity pattern was visibly disrupted (***Figure 3***a) and 24h rhythmicity was only weakly detectable in the mean behavior profile (eJTK, p=4.5 × 10^−3^; Table 1). The power and amplitude of the mean time series were lower than Aligned-cycle (Table 1; Dunn test, p<0.05; CircaCompare, p<0.05), and the difference in activity between dark and light phases was also severely reduced in terms of both time active (linear mixed effects model; LMM, p=1 × 10^−4^) and distance moved (LMM, p=2 × 10^−4^; ***Figure 3***c). Although 13/36 individuals (36%) were rhythmic (eJTK, p<0.001), their phases could not be distinguished from a circular uniform distribution (Rayleigh test, p=0.8), indicating desynchronization. Behavioral rhythms were also visibly disrupted and quantitatively weaker in Off10-cycle compared in Aligned-cycle (***Figure 3***a, Table 1), including reduced power (Dunn test, p<0.05), amplitude (CircaCompare, p<0.05), and dark:light activity ratio (LMM p=0.013). The phase of the mean time series in Off10-cycle was also advanced by 1.5 h (CircaCompare, p=5 × 10^−5^). The reduced dark-light activity ratios during Off10-cycle and Off12-cycle were specifically due to increased activity during lights-on (LMM, p<0.05), with no difference in activity during lights-off (LMM, p>0.1).

**Table 1.**
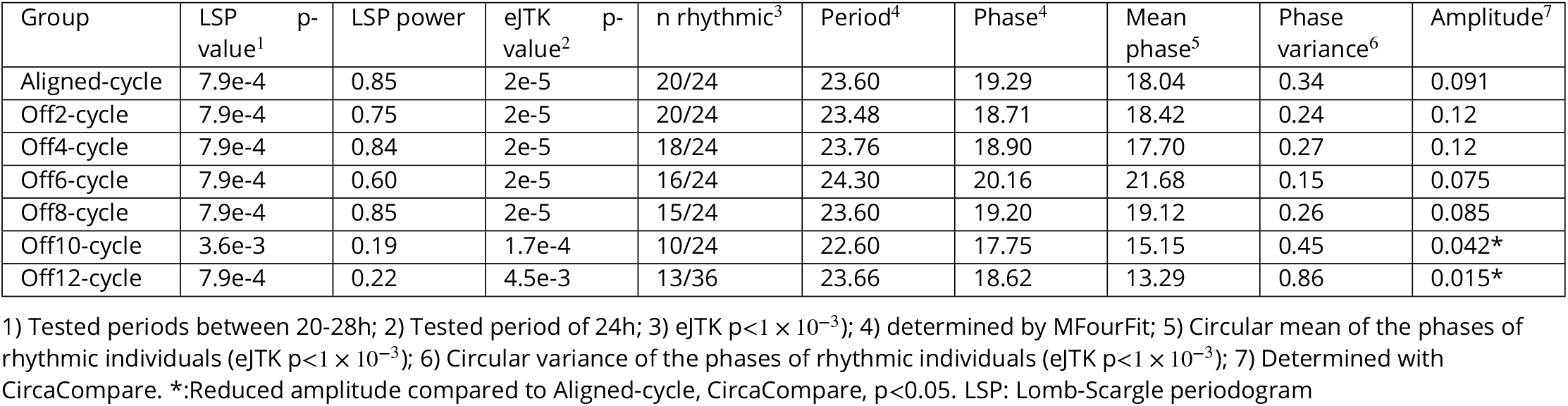
Summary statistics for sensory conflict experiments: **Cycling groups**

Weakened locotomor rhythms in Off10-cycle and Off12-cycle were caused both by a weakening of the rhythms of individual animals, and a lack of synchronization between the remaining rhythmic individuals. The percentage of rhythmic individuals declined monotonically with increasing SC, from 83% in Aligned-cycle to 36% in Off12-cycle (Table 1), and the LSP powers of animals in Off10-cycle and Off12-cycle were lower than during Aligned-cycle (Dunn test, *p* < 0.05; ***Figure 3–Figure Supplement 2***). In addition, the phases of rhythmic individuals in Off10-cycle and Off12-cycle had much larger variances than other groups and could not be distinguished from uniform distributions (Table 1; Rayleigh test, p>0.05). Thus, under severe SC, individual *Nematostella* were less likely to exhibit rhythmic behavior, and less likely for that behavior to synchronize with other individuals. Anemones exhibited robust diel behavior under smaller degrees of misalignment (0-8h; ***Figure 3***). The Off2-cycle, Off4-cycle, and Off8-cycle groups exhibited rhythms similar to those in Aligned-cycle, with no significant phase shifts or reductions in rhythm strength. The Off6-cycle group also showed 24h rhythms, but the mean phase of rhythmic individuals was delayed by 3.6h relative to Aligned-cycle (Watson test, p=0.0046); the mean behavior profile was also delayed (CircaCompare, p=4 × 10^−5^). The dark:light activity ratio was reduced (LMM, p=0.024) in Off6-cycle, and there was a marginally significant circatidal (10-14h) component (LSP, p=0.006). We confirmed that *Nematostella* exhibited normal diel behavior under a broad range of zeitgeber regimes using clustering analysis. We quantified distances between the 14 mean time series based on their spectral properties using wavelet transformation followed by principal component analysis and hierarchical clustering (see Methods). The Aligned-cycle, Off2-cycle, Off4-cycle, Off6-cycle, and Off8-cycle groups formed a single cluster with high support based on the approximately unbiased (AU) test, corresponding to normal rhythmic behavior, whereas Off10-cycle and Off12-cycle fell into different clusters (***Figure 4***). *Nematostella*’s diel behavior was therefore robust up to an 8h misalignment between light and temperature, although there was a detectable phase delay specifically in Off6-cycle.

**Figure 4.**
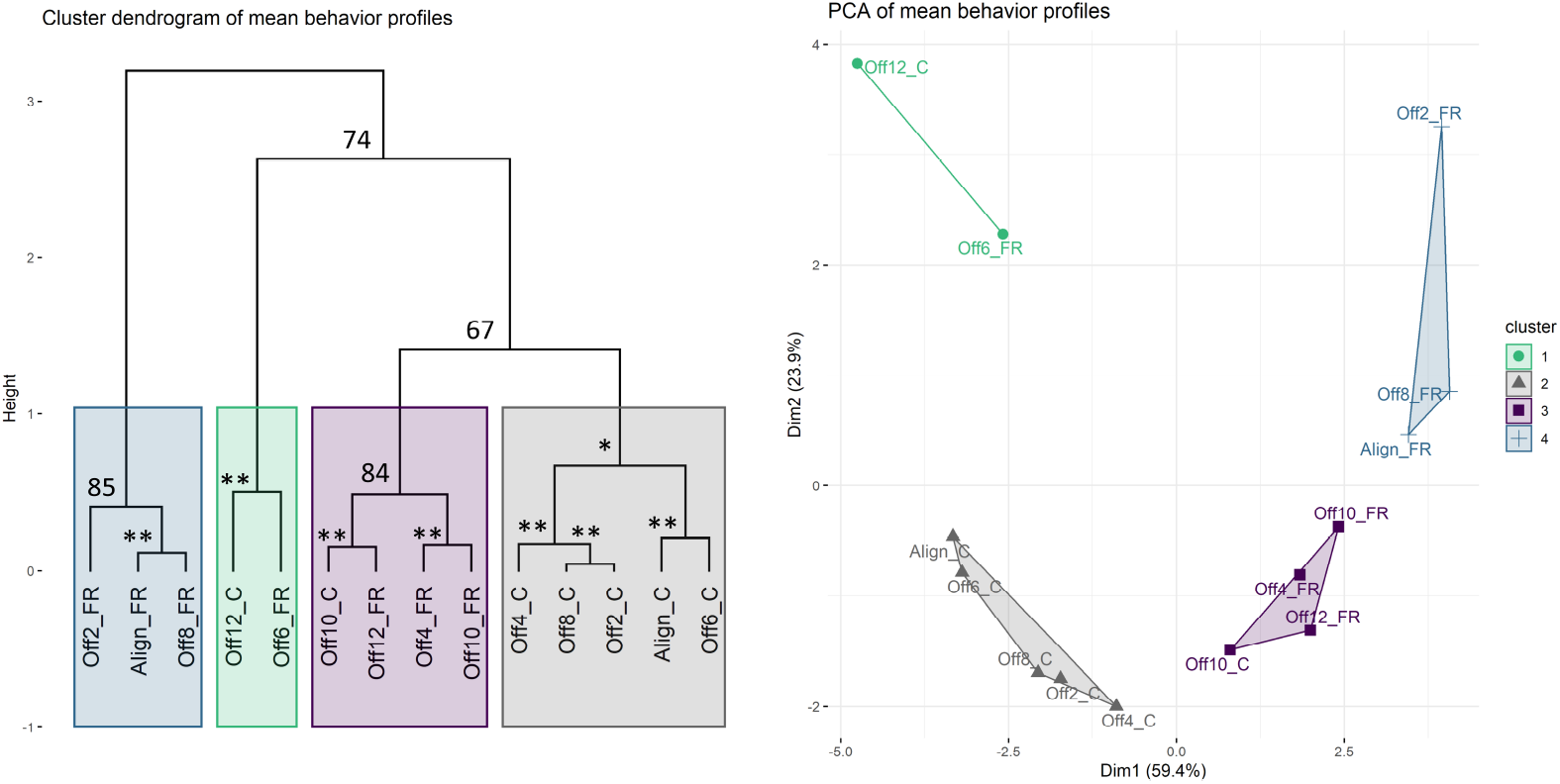
Clustering analysis of mean behavior profiles based on wavelet transformation. A) Principal components analysis of wavelet transformation distance matrix. Time series were grouped and colored based on the clusters identified in B). C: cycles; FR: free-running. B) Hierarchical clustering of samples in principal components space. Colored rectangles indicate the four clusters discussed in the text. Numbers indicate unbiased (AU) P values. *unbiased (AU) P value >= 90; **unbiased (AU) P value >=95.

### Sensory conflict disrupts endogenous rhythms

The above disruptions of behavior during SC could be due to masking effects rather than disruption of endogenous rhythms. However, we found that free-running behavior was also disrupted by SC, demonstrating *bona fide* effects on the endogenous clock (Tables 1 and 2, ***Figure 3***a). The mean behavior of animals in Off6-FR and Off10-FR was totally arrhythmic (LSP and eJTK, *p* > 0.1). Mean behavior in Off12-FR was rhythmic (eJTK, p=7 × 10^−5^), but the power and amplitude of the mean time series were reduced compared to Aligned-FR (Dunn test, p<0.05; CircaCompare, p<0.05), there were fewer rhythmic individuals, and the variance of the phases of rhythmic individuals was larger (Table 2). In the clustering analysis, Aligned-FR, Off2-FR, and Off8-FR formed one cluster with moderate AU support, while Off4-FR, Off10-FR, and Off12-FR formed a cluster that also included Off10-cycle (***Figure 4***). The final cluster was strongly supported and consisted of Off6-FR and Off12-cycle, two groups with severely disrupted rhythms.

**Table 2.**
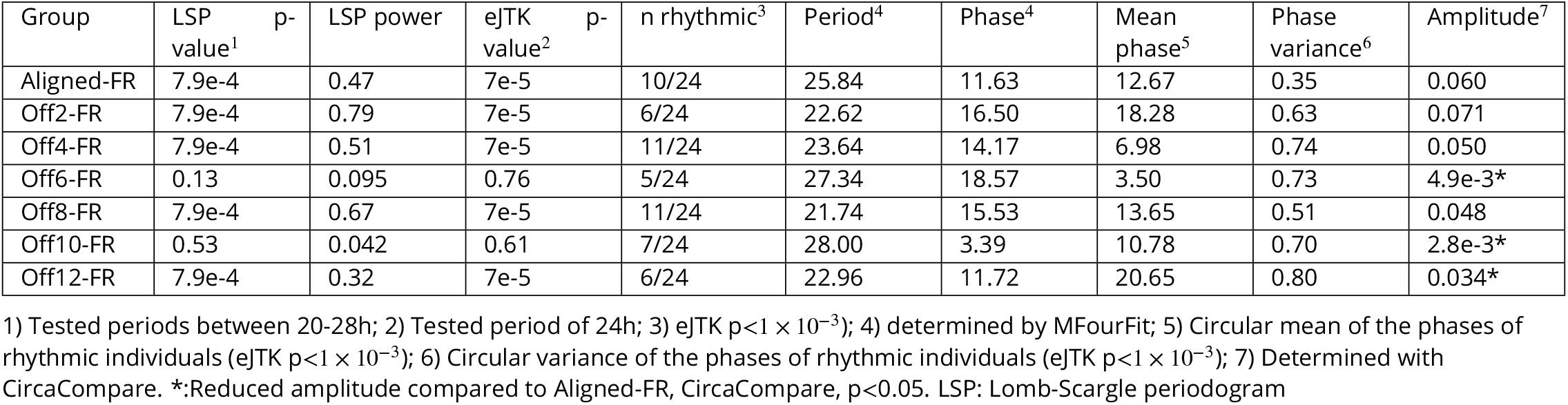
Summary statistics for sensory conflict experiments: **Free-running groups**

Behavioral patterns during free-running differed from patterns during cycling conditions. In fact, Off12-FR was more strongly rhythmic than Off12-cycle (Table 2). Thus, antiphasic zeitgeber cycles weakened, but did not abolish, endogenous circadian rhythms, and behavior was further disrupted during cycling conditions. Conversely, while endogenous rhythms were severely disrupted during Off6-FR, light and temperature cycles were able to drive rhythmic behavior during Off6-cycle (albeit with quantitative differences from Aligned-cycle). Additionally, disruptions of free-running rhythms did not increase linearly with the degree of offset. Behavior in Off6-FR was arrhythmic, but Off4-FR and Off8-FR exhibited clear rhythmicity (Table 2).

Disruption of free-running behavior was due primarily to a lack of synchronization among rhythmic animals, rather than weakened rhythms of individual animals. Only Aligned-FR had a significant non-uniform phase distribution (Rayleigh test, p<0.05). Mean powers of behavioral rhythms did not differ across free-running groups (Dunn test, p>0.05), nor was there a clear trend in the number of rhythmic individuals (Table 2). Phases of free-running rhythms differed in some groups, but not according to an obvious pattern. The phase of Off2-FR was advanced by 1.8h relative to Aligned-FR, Off8-FR was advanced by 4.4h, and Off12-FR was advanced by 4.3h (CircaCompare, p<7 × 10^−3^). Period length was not affected by SC (***Figure 3–Figure Supplement 1***).

### Relative strengths of light and temperature zeitgebers

To gain insight into the relative strengths of the zeitgebers, we compared the phases of rhythmic individuals against the offset between light and temperature (***Figure 3***d). In isolation, light and temperature each entrain rhythms that peak at roughly ZT18 (***Hendricks et al., 2012; Oren et al., 2015***); (***Figure 2***a), which we plot as the black and grey lines in ***Figure 3***d), respectively. If the phase of a rhythm were determined entirely by light, it would be close to the black line; if determined by temperature, it would be close to the grey lines. The phases of Off4-cycle and Off8-cycle groups were significantly closer to the light-entrained (black) line (paired Wilcoxon rank tests, p<0.05). All other groups were approximately equidistant between zeitgebers, including every free-running group. This illustrates that light exerts strong direct control on behavior, although light and temperature interact to set the phase of circadian behavior. When zeitgebers were close in phase (Aligned and Off2), they acted together the set the phase of locomotor rhythms, while at larger offsets phases were either determined primarily by the light cycle, or were intermediate between light and temperature. On the other hand, we found no evidence that the phases of free-running rhythms were primarily determined by either zeitgeber, with the caveat that this test had less power for free-running groups due to the numbers of rhythmic animals. Thus, although light has strong direct effects on behavior, it is not necessarily a stronger zeitgeber than temperature.

### Sensory conflict alters transcriptome-wide patterns of rhythmic gene expression

We used 3’ tag-based RNA sequencing (Tag-seq) to characterize the effects of SC on gene expression. Anemones were sampled at 13 time points across 48 h, either under Aligned-cycle or Off12-cycle conditions (***Figure 1***c). Reads were mapped to the SimRbase genome (Nvec200_v1, https://genomes.stowers.org/starletseaanemone).

SC substantially altered, and in many cases inverted, diel gene expression patterns in *Nematostella*. Under Aligned conditions, 1009 genes (6.6% of the transcriptome) were differentially expressed (DE) between light and dark samples (p<0.05), while 627 genes were DE between light and dark during SC, and 277 genes were DE in both time series. The correlation of log-fold changes of the 277 shared genes was negative (Pearson’s r=−0.59), meaning that genes upregulated in the light under Aligned conditions were generally downregulated in light under SC, and vice versa. Consequently, only 25 genes were DE between light and dark samples when averaged across both time series, and 1747 genes differed in their response to light across conditions. Using discriminant analysis of principal components (DAPC), we confirmed that genes distinguishing light and dark samples under Aligned conditions failed to do so during SC (***Figure 5***a). When light and dark SC samples were plotted on the discrimnant axis that maximized the variance between light and dark Aligned samples, the light SC animals were closer to the dark Aligned animals, showing that the diel expression of many genes was reversed.

**Figure 5.**
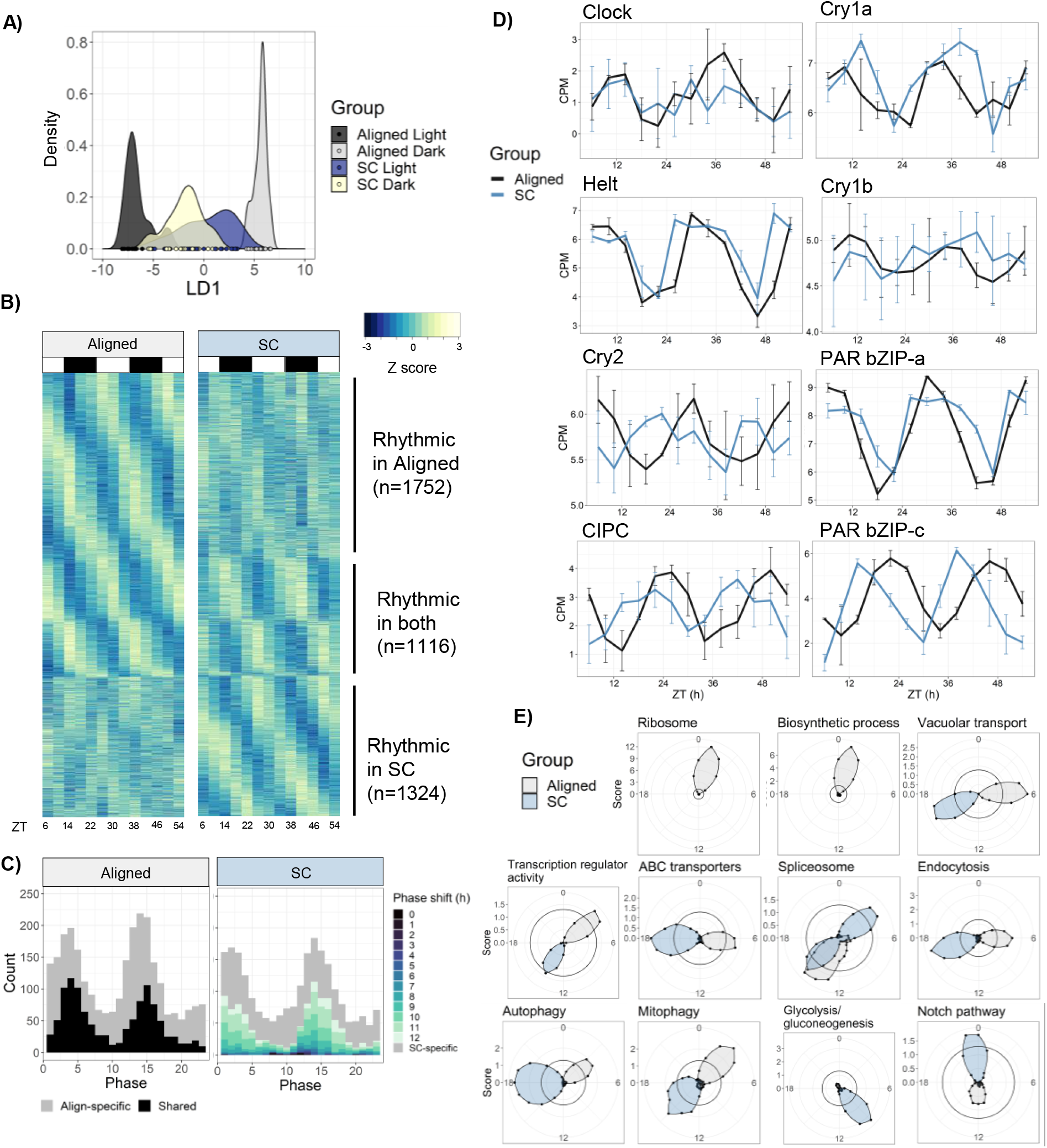
Sensory conflict alters patterns of rhythmic gene expression. **A)** Discriminant analysis of principal components (DAPC) demonstrates shifts in diel gene expression under SC. Density plot of sample loading values of light and dark samples during Aligned and SC conditions, on 1st discriminant axis from DAPC of aligned samples (“LD1”). **B)** Heatmaps of normalized rhythmic gene expression. Replicates were averaged at each time point and Z-scores were calculated for each gene and time series. “Rhythmic” genes had a RAIN p-value < 0.01 in the Aligned time series (top), SC time series (bottom), or both (middle). White and black rectangles represent light and dark time points, respectively. **B)** Phase distributions of rhythmic genes in Aligned (left) and SC (right) time series. Grey genes were rhythmic in only one time series, and black and colored genes were rhythmic in both. In SC, the color of shared rhythmic genes represents the phase shift of that gene from the Aligned time series. **D)** Phases of core clock genes under Aligned and SC conditions. Lines show mean counts per million, and error bars represent 95% confidence intervals. **E)** Sliding window enrichment analysis of select GO and KEGG terms during Aligned and SC time series. P-values were calculated by comparing genes with peak phase within a 4h sliding window with all other genes, and the score (y-axis) at each point is the average adjusted -log10 p-value at that time point (see Methods). Black circles indicates an FDR threshold of 0.05. **Figure 5–source data 1**. Counts matrix. Contains TMM-normalized counts-per-million on a log2 scale. **Figure 5–source data 2**. Differential expression, rhythmicity, and WGCNA analyses, and gene annotations. **Figure 5–source data 3**. Results of HOMER motif enrichment analyses. **Figure 5–source data 4**. GO and KEGG enrichment results.

Temperature was a stronger regulator of gene expression than light in our experiment: 1978 genes (13%) had a significant linear response to temperature, compared to only 25 DE genes in response to light. However, light-responsive genes notably included three putative circadian transcription factors (*PAR-bZIP-a, PAR-bZIP-c*, and *Helt*). Temperature-responsive genes were enriched for gene ontology (GO) terms related to transcription factor binding and oxidoreductase activity; genes with a significant Condition:Light interaction were also enriched for terms related to transcription factor activity. This suggests that the temperature cycle had widespread effects on transcription during light-dark cycles and modulated the expression of transcription factors. The mean expression level of 90 genes significantly differed between Aligned and SC time series. These genes were not enriched for any GO terms, but included one of the aforementioned PAR-bZIP proteins and a cytochrome P450 oxidase (CYP) previously implicated in circadian rhythms in *Nematostella* (***Oren et al., 2015***), along with two other CYPs and a SOX-family transcription factor (***Figure 5–source data 2***). Many genes (1898) displayed a linear change in expression with time, presumably because animals were not fed during the experiment. Genes with decreasing expression over time were enriched for GO terms related to cellular metabolism, suggesting an overall downregulation of metabolic processes as animals went without food. For complete lists of DE genes and GO enrichment results, see ***Figure 5–source data 2*** and ***Figure 5–source data 4***.

We searched 1kb upstream of transcription start sites (“promoters”) for putative binding motifs that may play a role in regulating circadian gene expression using the HOMER motif discovery tool. Genes whose mean expression during SC differed from mean expression during Aligned conditions were enriched for a motif of unknown function, which was most similar to Myb-related binding motifs in plants (HOMER, p=1 × 10^−12^; ***Figure 5–source data 3***), while genes with a negative Condition:Light interaction (i.e. genes with relatively more dark expression during SC) were strongly enriched for numerous bZIP transcription factor binding motifs (HOMER, p=1 × 10^−14^). Genes whose expression was negatively associated with temperature, and genes whose expression was positively associated with time, were also enriched for bZIP motifs (***Figure 5–source data 3***).

### Rhythmicity analysis

We tested for rhythmic gene expression using RAIN (rhythmicity analysis incorporating non-parametric methods) v1.24.0 (***Thaben and Westermark, 2014***), a non-parametric approach particularly suited for the detection of asymmetric waveforms. We compared RAIN to two other pieces of software: ECHO (extended circadian harmonic oscillator, (***De los Santos et al., 2020***), a parametric method based on the harmonic oscillator equation, and DryR (***Weger et al., 2021***), another parametric approach based on harmonic regression. Encouragingly, there was strong agreement between the three programs despite their completely different statistical methodologies (see ***Appendix 1***). We focus on the output from RAIN, and the output of the other two programs is available in ***Figure 5–source data 2***.

Surprisingly, there were similar overall numbers of rhythmic genes in Aligned and SC conditions, although less than half of them were rhythmic in both time series. Under Aligned conditions, 2868 genes displayed 24h rhythms in expression (RAIN, p<0.01; 18% of the transcriptome), while 2440 genes were rhythmic during SC, and 1116 genes were shared (***Figure 5***b). To define condition-specific rhythmic genes, we excluded genes that were marginally significant in the other condition (p<0.1); by this metric, 1225 genes lost rhythmicity during SC (Align-specific) and 921 gained rhythmicity (SC-specific). The promoters of shared rhythmic genes were enriched for bZIP transcription factor binding sites (p=1 × 10^−6^), while the promoters of Align-specific and SC-specific rhythmic genes were not enriched for any motifs (***Figure 5–source data 3***).

To understand how functional categories of rhythmic genes were affected by SC, we used a sliding window approach for functional enrichment of rhythmic genes (see Methods). This revealed dramatic changes to the expression of genes that mediate macromolecule, protein, and RNA metabolism during SC (***Figure 5***e). Most terms that were enriched among aligned rhythmic genes were not enriched at any time during SC, and only a single GO or KEGG term, “Spliceosome”, was enriched at the same time of day in both groups. Under aligned conditions, terms related to macromolecule and protein metabolism were enriched in early morning (ZT0-3) and lost enrichment under SC, while “RNA transport” and “ubiquitin-mediated proteolysis” were enriched in early night (ZT13-14) and lost rhythmicity under SC. Other terms shifted from day to night during SC, including “transcription factor activity”, “vacuolar transport”, “hydrolase activity”, “autophagy”, “mitophagy”, “ABC transporters”, and “mRNA surveillance pathway”; “protein processing in ER” shifted from ZT10 to ZT4 (***Figure 5–source data 4***). Some terms were only enriched under SC, including “DNA binding”, “Notch signalling pathway”, “ATP binding”, “GTPase activity”, “Glycolysis”, and “Pyruvate metabolism” (***Figure 5–source data 4***). For full enrichment results, see ***Figure 5–source data 4***. The phase distribution of Aligned rhythmic genes was bimodal (Rayleigh test, p=1 × 10^−4^), with peaks in mid-morning (ZT4) and early night (ZT14) (***Figure 5***c). Genes with phases during the two peaks (ZT0-6 and ZT12-18) were less likely to lose rhythmicity during SC (38% lost rhythmicity compared to 53% of other genes; Chi-squared test, p<2 × 10^−16^), suggesting that rhythmic gene expression during these windows was somehow more robust. The phase distribution under SC was also bimodal (***Figure 5***c), although the two distributions were not identical (Rao’s test for circular homogeneity, p=0.046). The vast majority of genes that were rhythmic under both Aligned and SC conditions (1092/1116, 98%) shifted in phase relative to the light cycle, with a median phase shift of 10.3h (CircaCompare, p<0.01); 86% shifted by at least 6h, and 58% by 10-12 h. Only 24 genes did not shift in phase relative to the light cycle, including *Clock, Helt*, and *PAR-bZIP-a* (***Figure 5***b). We identified 647 genes that shifted in phase by 10-12h relative to the light cycle and did not shift in phase relative to the temperature cycle; these genes closely tracked the phase of the temperature cycle and may be directly regulated by temperature. They were enriched for GO terms related to transcription factor activity, and the KEGG terms “Mitophagy” and “Spliceosome” (***Figure 5–source data 4***).

The amplitude and mean expression of rhythmic genes were slightly lower overall during SC. Mean expression of SC-specific genes was 18% lower than that of Align-specific genes during their respective time series (Wilcoxon, p=1 × 10^−8^; ***Figure 5–Figure Supplement 1***), but expression levels of shared rhythmic genes did not differ during SC (p=0.9). The amplitudes of Align-specific and SC-specific genes did not differ (p=0.49), but the amplitudes of shared rhythmic genes (n=1116) were median 7.1% lower during SC (Wilcoxon, p=2 × 10^−4^; ***Figure 5–Figure Supplement 1***). Specifically, amplitudes of the 647 temperature-responsive genes were reduced by median 8.8% (Wilcoxon, p=7 × 10^−4^).

### Comparison with light cycles at constant temperature

To assess how rhythmic gene expression during simultaneous light and temperature cycles compares to gene expression during a light cycle at constant temperature, we re-analyzed a previous study that sampled *Nematostella* during an LD cycle at 23°C (***Oren et al., 2015***). The phases of clock genes between (***Oren et al., 2015***) and the current study are shown in Table 3, and this comparison is discussed in more detail in ***Appendix 2***. The phases of core circadian genes (*Clock, Helt, CIPC*, cryptochromes, and PAR-bZIPs) were within 4h across our Aligned time series and in (***Oren et al., 2015***), with the exception of *Cry1b*. However, there was little overlap between rhythmic genes overall, as only 477 / 1498 (32%) genes rhythmic in Oren et al. (RAIN, p<0.05) had a p-value less than 0.05 in the current study. Differences in clock output may be due to experimental conditions or differences between populations. Nonetheless, our re-analysis shows that core clock gene expression was not altered by the addition of a temperature cycle in-phase with the light cycle.

**Table 3.** Comparison of phases of putative clock genes

| Study | Clock | Helt | CIPC | PAR-bZIP-a | PAR-bZIP-c | Cry1a | Cry1b | Cry2 |
| --- | --- | --- | --- | --- | --- | --- | --- | --- |
| Current study (Aligned) | 12.4 | 9.2 | 1.0 | 7.3 | 22.3 | 9.3 | 10.5 | 5.6 |
| Current study (SC) | 11.94 | 7.9 | 19.3 | 8.0 | 16.2 | 12.1 | NA | 21.8 |
| Oren et al. (2015)* | 10.9 | 7.3 | 23.4 | 5.3 | 19.4 | 7.9 | NA | 2.3 |
| SimrBase ID | 6258 | 15017 | 4651 | 8136 | 8448 | 8109 | 8041 | 15282 |
| JGI ID | 160110 | 246249 | 245026 | 150375/87565 | 39846 | 168581 | 106062 | 194898 |

Phases estimated with CircaCompare. NA: below significance cutoff, phase not calculated. We used a p-value cutoff of $p = 0.01$ for the current study, and $p = 0.05$ for (Oren et al., 2015) because of their small sample size.

### Network analysis

We used weighted gene co-expression network analysis (WGCNA) to characterize co-expressed modules of rhythmic genes. First, to understand how SC changed the rhythmic transcriptome, we compared networks constructed separately for genes rhythmic in each time series (Align-specific and SC-specific). Second, we constructed a Full network using all genes rhythmic in either time series. The timing of module expression in the Full network is shown in ***Figure 6***b, the expression of select modules is shown in ***Figure 6***a, and the relationships between modules in all three networks are shown in ***Figure 6***c. Functional enrichment results are available in ***Figure 5–source data 4***. Modules are referred to with color names followed by an abbreviation indicating their network. For example, the Blue module in the SC-specific network is Blue-SC, whereas the Blue module in the Full network in Blue-F.

**Figure 6.**
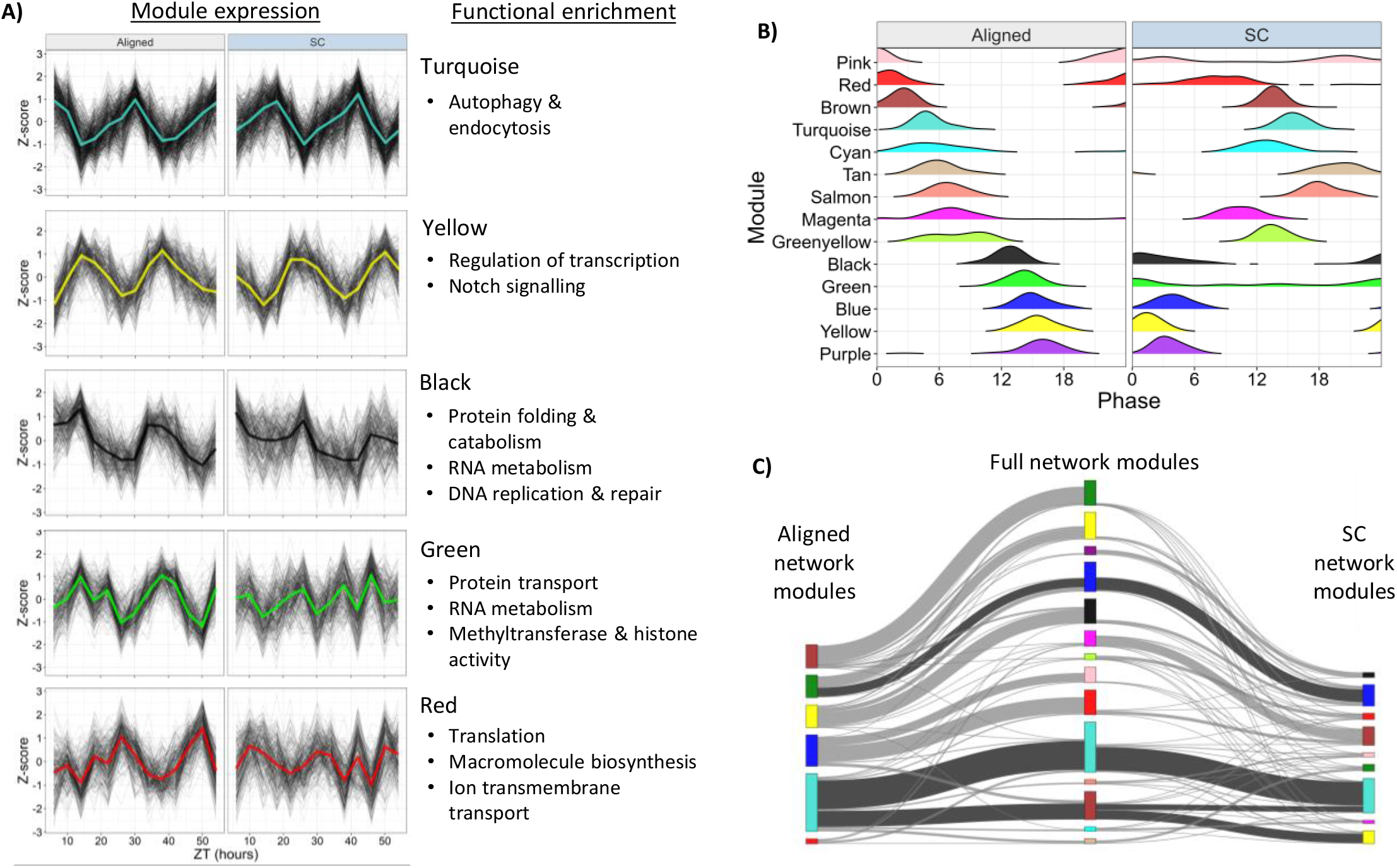
Network analysis identifies changes to rhythmically co-expressed gene modules. **A)** Expression patterns of select modules in the full network. Black lines are Z-scores (difference from mean divided by standard deviation) of the expression of each gene, and the colored line is the mean of those Z-scores. Left panels are Aligned samples, and right panels are SC samples. Functional enrichment **B)** Phase distributions of modules genes in the Full network during Aligned (left) and SC (right) conditions. Each line shows a smoothed density estimate of the phases of rhythmic genes within that module, weighted by the -log10(p-value) from RAIN. **C)** Sankey plot showing gene overlap between modules in the Align-specific (left), Full (middle), and SC-specific (right) networks. Height of module bars represents the number of genes in that module, and the width of connections represents the number of shared genes. Dark connecting lines indicate highly significant (p<1 × 10^−5^) overlap between Aligned and SC network modules. Genes in grey modules are not shown. Color labels were assigned independently for each network (no inherent meaning).

Genes in the Align-specific network (2868 genes, 39 samples) were assigned to 6 modules, with 889 genes (31%) unassigned. All module eigengenes—the first principal component of the module expression matrix—were rhythmic (LSP, p<0.01). This network provides a picture of “normal” rhythmic gene expression, and functional enrichment recapitulated many of the results from our sliding window analysis. Modules with daytime peak expression were enriched for processes related to translation, ribosomes, and nitrogen compound biosynthesis (Blue-A); and transcription factor activity, vacuolar transport, and GTPase activity (Turquoise-A). A module with peak expression at ZT12 was enriched for catabolic processes, including proteolysis and RNA degradation (Yellow-A). Modules with night-time peak expression were enriched for ATP binding, chaperone activity, chromatin organization, and the spliceosome (Brown-A); and “negative regulation of cellular macromolecule biosynthetic process” (Green-A). Genes in the SC-specific network (2440 genes, 39 samples) were assigned to 9 modules, with 1080 genes (44%) unassigned. Most genes in the Aligned network were either arrhythmic or unassigned in the SC network (***Figure 6–Figure Supplement 1***). SC modules with peak daytime expression were enriched for a single GO term, “ion channel complex” (Black-SC); and the KEGG terms “Spliceosome” and “Protein processing in the ER” (Blue-SC). A module with peak expression in the late day was enriched for “protein folding” and several KEGG terms related to carbon and amino acid metabolism, including “Glycolysis” (Brown-SC). Modules with peak night-time expression were enriched for proteolysis and peptidase activity (Pink-SC); terms related to DNA repair (Red-SC); and transcription factor activity, endocytosis, mitophagy, and autophagy (Turquoise-SC). Overall, just 25% of GO and KEGG terms enriched among Aligned network modules were enriched in any SC network module. This confirms that SC substantially disrupted the rhythmic expression of many functionally-related groups of genes.

However, some processes maintained coordinated expression during SC. Two modules (Turquoise-A and Green-A) strongly overlapped with modules in the SC network (hypergeometric test, p<<1 × 10^−5^; ***Figure 6***c) and two other modules had moderate overlap (Yellow-A and Red-A, p<0.01). Turquoise-A shared functional enrichment related to autophagy, endocytosis, and transcription factor activity with Turquoise-SC and Yellow-SC, indicating that these genes shifted in phase concordantly during SC.

To understand how SC affected higher-order network properties, we quantified the preservation of underlying network characteristics during SC using module preservation statistics (see Methods and ***Langfelder et al., 2011***, for details). These statistics test whether the density and connectivity of genes in Aligned network modules are higher in the SC network than between random pairs of genes from the SC network. Density and connectivity were broadly conserved across all six Aligned modules and in a random sub-sample of genes from the Aligned network (“Gold” module; Table 4)—this is probably because many genes shifted in phase by 12h during SC, maintaining similar co-expression. However, although SC did not totally disrupt co-expression patterns, they were substantially weakened. Intramodular connectivity (kIM) was 43% lower in the SC network, and the connectivity of each gene to all other genes in the network (kTotal) was 31% lower (Wilcoxon, p<1 × 10^−15^). In the Full network (4192 genes, 78 samples), which assigned rhythmic genes to 14 modules with 932 genes (22%) unassigned, kIM was reduced by 23% during SC, and kTotal by 12% (Wilcoxon, p<1 × 10^−15^). The correlation between genes and their respective eigengene (kME) was reduced by 5.6% (Wilcoxon, p=6 × 10^−15^), and the proportion of each gene’s expression variance explained by eigengene expression (propVar) was reduced by 8.2% (Wilcoxon, p=1 × 10^−11^). kME, kIM, and propVar were significantly lower during SC whether they were calculated for all genes assigned to a module (n=3260), only for assigned genes rhythmic in their respective time series (n=2417 and n=1893), or only for assigned genes rhythmic in both time series (n=1050). By these metrics, some modules gained denser expression during SC (Blue-F, Purple-F, Magenta-F, and Greenyellow-F; n=722), but they contained fewer than half as many genes as modules that became less densely-expressed (Turquoise-F, Red-F, Black-F, Green-F, and Pink-F; n=1702). Together, these results show that rhythmic gene expression was less well-correlated both within and across modules during SC.

**Table 4.** Preservation statistics of modules identified in the Aligned network, assessed in the SC network

| Module | Z-summary | Z-density | Z-connectivity |
| --- | --- | --- | --- |
| Gold | 22.3 | 13.4 | 31.1 |
| Grey | 9.8 | 4.9 | 14.8 |
| Blue | 24.2 | 29.0 | 19.4 |
| Brown | 26.5 | 33.0 | 20.1 |
| Green | 18.9 | 27.8 | 10.1 |
| Red | 9.7 | 13.8 | 5.6 |
| Turquoise | 34.4 | 49.8 | 19.0 |
| Yellow | 16.8 | 22.7 | 11.0 |

The “Gold” module is a random sub-sample of 1000 genes from the Aligned network; the “Grey” module is rhythmic genes that were not assigned to any module in the Aligned network.

Analysis of the Full network illustrated that temperature was the most important driver of rhythmic gene expression. Half (7/14) of module eigengenes in the Full network maintained strong rhythmicity during SC (LSP, p<0.01), and all shifted in phase by 10-12h, following the temperature cycle. The Turquoise-F module (n=614) was enriched for terms related to autophagy and endocytosis, and the Brown-F module (n=341) was enriched for transcription factor activity. Turquoise-F and Brown-F promoters were enriched for bZIP binding motifs (HOMER, p<1 × 10^−8^), and both modules were negatively correlated with temperature (linear model, p<0.05). The Tan-F module (n=61) was enriched for ATP hydrolysis and ABC transporter activity, and the Salmon-F module (n=55) was enriched for serine-type endopeptidase activity; its promoters were enriched for ETS binding motifs (HOMER, p=1 × 10^−5^). The Black-F (n=298), Blue-F (n=359), and Yellow-F (n=327) modules were positively correlated with temperature (LM, p<0.05). Black-F was enriched for terms related to protein and RNA metabolism, chromatin organization, and DNA replication and repair. Blue-F was enriched for “Spliceosome”, “protein processing in ER”, and “Butanoate metabolism.” Yellow-F was enriched for regulation of transcription, protein dimerization activity, and Notch signalling. The Green-F (n=302) and Pink-F (n=189) eigengenes lost rhythmicity during SC. Green-F was enriched for terms related to RNA metabolism, protein transport, and regulation of gene expression, and contained the *Clock* gene. Green-F module promoters were enriched for putative binding motifs with unknown function (HOMER, p=1 × 10^−10^). The Pink-F module (n=189) was enriched for “Sphingolipid metabolism”. The Red-F module (n=299) was marginally rhythmic during SC (LSP, p=0.019), and advanced in phase by 8h (***Figure 6***b). This module was enriched for proton transmembrane transport and terms related to translation. Two module eigengenes gained rhythmicity under SC. Purple-F (n=101) was enriched for thiamine and purine metabolism, and its promoters were enriched for helix-turn-helix (HTH) binding motifs (HOMER, p=1 × 10^−5^), while Greenyellow-F (n=74) was not enriched for any functional terms. The Cyan-F (n=52) and Magenta-F (n=188) modules were marginally rhythmic in Aligned conditions (LSP, p=0.02), and shifted in phase by 4h and 8h, respectively, during SC. Cyan-F was enriched for the KEGG terms “Glycolysis” and “TCA cycle”, but this enrichment was driven only by two genes, both of which were annotated as phosphoenolpyruvate carboxykinases (PEPCK) (***Figure 5–source data 2***); Cyan-F promoters were enriched for SOX binding motifs (HOMER, p=1 × 10^−10^). Magenta-F was enriched for terms related to protein catabolism, oxidoreductase activity, and RNA transport.

## Discussion

We show here that ecologically relevant temperature cycles drive and entrain circadian behavior in the cnidarian *Nematostella vectensis*, and explore how the relationship between simultaneous light and temperature cycles affects rhythmic behavior and gene expression. Given that white light exerts strong direct effects on behavior in *Nematostella*, including masking free-running locomotor rhythms (***Oren et al., 2015; Tarrant et al., 2019***), we thought behavior might simply synchronize to the light cycle to the exclusion of temperature, as is the case for *Drosophila* peripheral clocks (***Harper et al., 2017***). However, when zeitgeber phases differed by 10-12h, *Nematostella*’s behavior became severely disrupted and arrhythmic on average. Free-running rhythms were also disrupted, depending on the specific phase relationship. This shows that both light and temperature interact to set the phase of *Nematostella*’s clock, and that neither cue dominates the other. As a consequence, normal rhythmic behavior is only possible under certain relationships between zeitgebers. The disruptive effects of sensory conflict (SC) were not acute responses to transient conditions, but chronic changes measured after weeks of exposure and acclimation.

Gene expression rhythms were also dramatically altered by SC. Surprisingly few genes directly responded to light, whereas several hundred genes followed the temperature cycle. Rather than simply reducing the number of rhythmic transcripts, SC caused turnover in the identities of rhythmic genes and altered the relative timing of expression between them. This resulted in widespread perturbation of the temporal expression of metabolic processes and an overall weakening of rhythmic co-expression patterns. Nonetheless, many genes were robustly rhythmic during SC, and other genes even gained rhythmic expression. These patterns differed between functional categories. For instance, genes related to autophagy and endocytosis shifted in phase following the temperature cycle, while genes related to ribosome and protein metabolism lost rhythmicity.

### Simultaneous light and temperature cycles produce complex behavior

Rhythmic behavior is the net result of endogenous and exogenous signals, leading to potentially complex behavioral outcomes. Direct effects of environmental cues can mask endogenous rhythms (***Aschoff, 1960***), and, conversely, circadian rhythms can constrain or “gate” acute environmental responses (***Salter et al., 2003; Fowler et al., 2005***). During simultaneous light and temperature cycles, we found that behavior was increasingly disrupted up to the largest degrees of misalignment (10h and 12h offsets), and there was an underlying weakening and disruption of endogenous rhythms. However, behavioral patterns during zeitgeber cycles did not directly mirror the underlying free-running behavior, indicating that direct effects of light and temperature interacted with and masked endogenous behavior. Our data illustrate two contrasting scenarios for the relationship between endogenous and exogenous cues. During the 6h offset, endogenous rhythms were strongly disrupted, but behavior during cycling conditions was roughly normal. In this case, light and temperature directly drove rhythmic behavior, although the underlying clock output was arrhythmic. On the other hand, during the 12h offset, endogenous rhythms were relatively robust, but behavior during cycling conditions was arrhythmic. In this case, misaligned light and temperature cycles masked endogenous rhythms.

SC provides a framework to perturb circadian systems and characterize the range of environmental conditions under which normal clock output is possible. Remarkably, differences of as little as two hours resulted in completely different behavioral outcomes, revealing non-linear dynamics in free-running behavior. Free-running rhythms were most strongly disrupted at intermediate (6h and 10h) offsets, and somewhat weakened at the 12h offset. However, rhythms were robust at the 4h and 8h offsets. The underlying reasons for these non-linearities probably depend on the molecular pathways that connect light and temperature to the clock, which are unknown. These findings highlight the need for multi-zeitgeber studies to employ fine temporal resolution rather than simply testing two alternative phase relationships. Similar experiments in other systems will likely reveal similarly complex behavioral outcomes, opening the door for comparative studies of clock sensory processing in animals.

### Implications for the cnidarian molecular clock

The molecular architecture of circadian clocks is understood in bilaterian model systems, but is largely unknown in Cnidaria. *Nematostella* possesses orthologs of the core circadian transcription factors *Clock* and *Cycle*, but not of the canonical repressor of CLOCK, *Period* (***Reitzel and Tarrant, 2009; Reitzel et al., 2013***). This suggests that there are major differences in clock architecture between bilaterians and cnidarians. We identified binding motifs enriched in the promoters of rhythmic genes, suggesting that bZIP, and possibly SOX, family transcription factors may regulate circadian gene expression in *Nematostella*. Proline and acidic amino acid-rich bZIP (PAR-bZIP) and SOX-family proteins regulate circadian clock outputs in bilaterians (***Cyran et al., 2003; Gachon, 2007; Cheng et al., 2019***), and *Nematostella* PAR-bZIP and SOX genes were robustly rhythmic during Aligned and SC conditions in our data (***Figure 5–source data 2***). Our results also support existing hypotheses about the roles of cryptochrome proteins in the *Nematostella* clock, and raise new questions about the role of the *Clock* gene in this animal.

Our data support the hypothesis that the cryptochrome protein *Cry2* is involved in the core clock, possibly as a negative regulator of *Clock* (***Reitzel et al., 2013***), while *Cry1a* and *Cry1b* are photosensors. *Cry2* expression was driven by both light and temperature cycles, while rhythmic expression of *Cry1a* and *Cry1b* appears to be driven only by light cycles. Other genes suspected to have central roles in the cnidarian clock, including *Helt-like* (*Helt*), *CIPC*, and two PAR-bZIP transcription factors (*PAR-bZIP-a* and *PAR-BZIP-c*) (***Reitzel et al., 2010; Oren et al., 2015; Tarrant et al., 2019***), were strongly rhythmic in our study. Misalignment between light and temperature cycles did not disrupt oscillations of these putative clock genes, as (except for *Cry1b*) they were all strongly rhythmic during both aligned and SC conditions. *CIPC, Cry2*, and *PAR-bZIP-c* advanced in phase by 5-8h under SC, while *Clock, Helt*, and *PAR-bZIP-a* remained in-phase with the light cycle. Thus, although temperature was a stronger driver of rhythmic gene expression than light overall, light probably controls rhythmic expression of some core clock components.

Surprisingly, we found that *Clock* mRNA did not oscillate during a temperature cycle (***Figure 2***), and was one of only two dozen genes to directly follow the light cycle. Our data, together with evidence that *Clock* expression is induced by blue light and not green light (***Leach and Reitzel, 2020***), suggests that rhythmic *Clock* expression is entirely blue light-driven and is not required for rhythmic behavior. This would be unusual, as we are not aware of any other system in which *Clock* mRNA oscillates in response to one zeitgeber and not another: *Clock* oscillates during both light and temperature cycles in *Drosophila* (***Glossop et al., 1999***) and fish (***Lahiri et al., 2005; Di Rosa et al., 2015***), and *Cycle/Bmal* does the same in mammals (***Chun et al., 2015***). Strictly speaking, we do not even know whether *Clock* is required for circadian rhythms in *Nematostella*, but it is possible that *Clock* exhibits temperature-driven rhythms in a subset of cells, or that temperature acts on *Clock* activity at a level other than transcription.

### Global patterns of gene expression during SC

To our knowledge, this is the first study of transcriptome-wide gene expression during SC in any organism. Because 12h SC disrupted rhythmic locomotion, we had expected that *Nematostella* might simply express fewer rhythmic genes in these conditions. Instead, while many genes did lose rhythmicity during SC, several hundred genes in turn became rhythmic, and 7% of the transcriptome remained rhythmic. Thus, a substantial portion of the transcriptome was rhythmic even during conditions that severely disrupted rhythmic behavior. It is unclear whether the cycling of genes that gained rhythmicity under SC is unimportant, deleterious, or serves some sort of homeostatic function. Gain or loss of rhythmic RNA abundance does not necessarily imply changes to rhythmic transcription, but could be due to rhythmic variation in splicing, degradation, or any other post-transcriptional process (***Lück et al., 2014; Zhang et al., 2014***). Indeed, we observed widespread changes to the expression of genes that regulate RNA metabolism and splicing during SC, possibly affecting the rhythmic abundance of many other transcripts. We also note that, although the amplitude of SC-specific rhythmic genes did not differ from Align-specific genes, they had lower mean expression. Nonetheless, it would be interesting to test whether SC can induce novel rhythmic expression at the protein level. The regulation of a large portion of the transcriptome by diel temperature cycles may be expected for ectothermic animals (***Boothroyd et al., 2007***), and some physiological processes in *Nematostella*, such as autophagy, may be robustly rhythmic because they are strongly synchronized by ambient temperature. However, the amplitude of temperature-responsive genes was reduced during SC, suggesting that some temperature-responsive genes are also either regulated by light, or by the clock itself. Furthermore, temperature-responsive genes were enriched for processes known to be under clock control in other animals, including autophagy, ABC transporter activity, and mRNA metabolism (***Ma et al., 2012; Hardin and Panda, 2013; Pácha et al., 2021***).

*Nematostella* experienced dramatic perturbations to metabolic gene expression during SC. For instance, genes that mediate ribosome biogenesis, one of the most energy-intensive cellular processes (***Warner, 1999***), lost rhythmicity, as did other genes related to RNA and protein metabolism. Desynchronization of metabolic processes likely has bioenergetic consequences, as circadian clocks normally optimize metabolism for periods of high and low energy demand (***Thurley et al., 2017***). Even genes that remained rhythmic and followed the temperature cycle, such as those related to endocytosis, vacuolar transport, and autophagy, changed their temporal expression in relation to time of feeding, and thus to the period of peak energy availability. Additionally, some genes related to glucose metabolism gained rhythmic expression during SC, suggesting that clock dysregulation altered the expression of glucose regulatory genes. In general, circadian clocks regulate metabolic homeostasis and receive input from metabolic pathways (***Ma et al., 2012; Rey et al., 2016; Hurley et al., 2018***). For this reason, disruption of clock function is linked to various metabolic diseases in mammalian models and human health (e.g. ***Turek et al., 2005; Green et al., 2008; Nedeltcheva and Scheer, 2014; Roenneberg and Merrow, 2016***). The desynchronization of rhythmically expressed metabolic gene modules by conflicting zeitgeber regimes plausibly contributes to negative organismal outcomes, and future work in *Nematostella* should connect these results to other physiological endpoints.

### What causes sensory conflict?

Although extreme conflict between zeitgebers may not reflect natural conditions, it is relevant to the artificial conditions that commonly perturb circadian rhythms in humans (***Roenneberg and Merrow, 2016***) and affect many other species via light pollution (***Gaston et al., 2013***). Generally, our results illustrate that disruption of circadian rhythms is not an all-or-nothing situation. Even when behavior was severely disrupted on average, some individuals were always able to maintain rhythmic behavior (although their rhythms were not synchronized with one another). SC did not abolish rhythmic expression of core clock genes, showing that clocks were functional on some level. Instead, SC caused clock outputs to become weaker and less coordinated among individuals. Investigating the determinants of individual variability in circadian output, including possible genetic factors, is a fascinating area of future research.

Disruptive effects of SC could be caused by desynchronization of multiple clocks in different tissues, or by disruption of individual clocks. In bilaterians, central and peripheral clocks can differ in their responses to SC, which might contribute to organism-level disruptions (***Harper et al., 2016, 2017***). On the other hand, dissociation of clock gene expression can occur even within single cells (***Schmal et al., 2019***). We found that SC disrupts behavior in an animal without a brain, meaning that desynchronization of central and peripheral clocks is not the only factor that contributes to disrupted rhythmic behavior. Nonetheless, *Nematostella* is not simple or homogeneous at the tissue level, possesses many different neuronal subtypes (***Sebé-Pedrós et al., 2018***), and may possess multiple, spatially distinct clocks. Future work to identify cellular clocks will help determine whether behavioral disruptions result from disruption of individual clocks, or desynchronization between multiple clocks distributed throughout the body.

## Materials and methods

### Animal culture

Animals used in this study were adult anemones of mixed sex. The laboratory population was originally collected from Great Sippewissett Marsh, MA USA and maintained in the laboratory for several generations. *Nematostella* were kept in glass water dishes containing half-strength 1 μm-filtered seawater (from Buzzards Bay, MA), diluted 1:1 with distilled water to a salinity of approximately 16. Water was changed weekly. Prior to acclimating to experimental conditions, anemones were kept at room temperature (18 °C) and fed brine shrimp nauplii four times per week.

### Behavioral experiments

Anemones were acclimated to a given light and temperature regime for at least two weeks prior to behavioral monitoring. They were fed during the acclimation period, but not during behavioral monitoring. A schematic of the experimental design is given in ***Figure 1***. For all experiments, animals were randomly selected from the full acclimated group.

In the first set of experiments, *Nematostella* were kept in gradually changing (ramped) 24h temperature cycles in constant darkness. Incubators were programmed such that the water temperature reached the desired temperature on the hour, which was monitored with HOBO temperature loggers. Temperature cycles were either 14–26 °C (changing 1 degree per hour) or 8–32 °C (2 degrees per hour). Animals were fed with the aid of a dim red-light headlamp (*Nematostella* behavior is most sensitive to blue light and least sensitive to red light (***Reitzel et al., 2010; Leach and Reitzel, 2020***). Animals were transferred to the Noldus chambers at ZT6 (14–26 °C), or ZT8-10 (8–32 °C). Trial lengths ranged from 69-73 hours. For the 14–26 °C cycle, the free-running temperature was 20 °C, which was held constant beginning at ZT6. For the 8–32 °C cycle, the free-running temperature was 24 °C, beginning at ZT8.

For sensory conflict (SC) experiments, animals were acclimated to the 14–26 °C temperature cycle and a 12:12 light-dark cycle. The population of anemones was maintained in an incubator with “aligned” light and temperature cycles, such that ZT0 corresponded to lights-on and the coldest temperature. In addition to Aligned-cycle, 6 light and temperature regimes were tested in which the phase of the temperature cycle was delayed relative to the light cycle in 2h increments, as shown in ***Figure 1***b. Anemones in these groups were transferred from the aligned incubator to a second incubator where the temperature cycle was shifted relative to the light cycle. Behavioral monitoring was performed for separate groups of animals during cycling and free-running conditions, resulting in 14 total groups each with at least n=24 anemones (Off12-cycle had n=36). Minimum sample size of 24 was chosen because it was twice the sample size (n=12) at which we regularly detect rhythms in LD.

After acclimation, behavior was monitored using Noldus Daniovision observation chambers equipped with infrared-sensitive cameras (DVOC-0040, Noldus Information Technology). Anemones were fed the day before each behavioral trial. A few hours after feeding, 12 anemones were transferred to individual wells of two 6-well plates and left in the incubator overnight. This was done to avoid directly handling the animals at the start of the experiment. At the start of each trial, plates were placed into two Noldus chambers. A custom flow-through water system was used to control the temperature. A separate tank was heated and cooled using an aquarium heater and cooler connected to a programmable temperature controller, and water was flowed around the plate inside the observation chamber using aquarium pumps. Water temperature was monitored with iButton temperature loggers. Each trial began at ZT6, and was recorded for 78 h. Free-running trials begain at ZT12, 6h after the start of recording, and thus had a length of 72 h. To avoid suddent jumps in temperature, it was necessary to release groups into constant temperature (20 °C) at different times (see ***Figure 1***b; the beginning of free-run *per se*, meaning lights-off and 20 °C, was the same for every group). Videos were recorded with a framerate of 2 frames per second.

### qRT-PCR

Anemones were maintained in a 8–32 °C temperature cycle in constant darkness and sampled every 4h over 48 h (13 time points), beginning at ZT12. In parallel, another group of animals was moved to constant temperature (24 °C) at ZT8 and sampled at the same time points. Four biological replicates were sampled at each time point, and each replicate consisted of three pooled individuals. Total RNA was extracted using the Aurum Total RNA Fatty and Fibrous Tissue Kit (Bio-Rad) following manufacturer’s protocol with DNAse treatment. RNA quality and concentration were checked with a NanoDrop spectrophotometer. Libraries with low 260/230 ratios (< 1.8) or low concentration were re-purified using an RNA Clean and Concentrator kit (Zymo). RNA libraries were diluted to 20 ng μL^−1^ and reverse transcription was performed with an iScript cDNA synthesis kit (Bio-Rad). Primers for five genes—*Clock, Cry1a, Cry1b, Cry2*, and *Helt*—were ordered from Thermo Fisher and diluted to 10 μM. Primer sequences are in Table 5. Quantitative PCR (qPCR) was performed with iTaq Universal SYBR Green Supermix (Bio-Rad). The 20 μL reaction mixture consisted of 10 μL Supermix, 8 μL nuclease-free water, 1 μL cDNA, and 0.5 μL each of forward and reverse primers. All samples for a given primer pair were performed on a single 96-well plate. Annealing temperatures were tested with a gradient for each primer pair followed by a melt curve analysis. 60 °C was chosen as the annealing temperature because it resulted in the lowest C_q_ values with specific amplification. The thermocycler was programmed as follows: 95 °C for 3 min, 40 cycles of 95 °C for 10 s and 60 °C for 30 s, followed by a melt curve analysis.

**Table 5.**
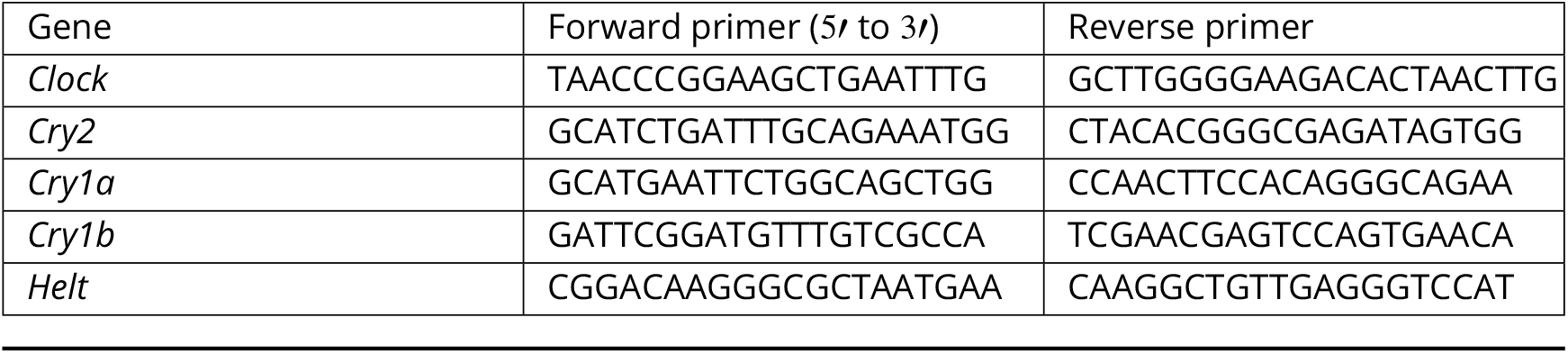
Primers used for qRT-PCR

Quantitative PCR data analysis was performed with LinRegPCR (***Ruijter et al., 2009***). We excluded a total of three samples from further analysis because several genes failed to amplify. Gene expression was normalized using the NORMA-Gene algorithm (***Heckmann et al., 2011***). Rhythmicity was tested with Lomb-Scargle periodograms (LSP) implemented in the R package ‘lomb’ v2.0, with periods between 20-28 h. Significance was assessed with n=2000 permutations, with a slight modification to the default ‘randlsp’ command. Since each time point contained replicates, all 49 data points (3-4 replicates x 13 time points) were shuffled, and then the mean time point values were re-calculated for the permuted data.

### RNA extraction and sequencing

Anemones were either maintained in Aligned-cycle or Off12-cycle conditions (***Figure 1***c). Animals were sampled every 4h over 48 h (13 time points), beginning at ZT6, and immediately preserved in RNA-later. Each time point contained three biological replicates of five pooled individuals each, with the exception of one library that consisted of three pooled individuals. This was because the initial 5-animal pool produced a low yield, so a new pool was constructed from 3 remaining animals from that treatment group. RNA was extracted using the Aurum Total RNA mini kit following the manufacturer’s instructions, except we incubated the RNA lysis buffer on ice for 30 min following tissue homogenization. Concentrations and quality were assessed with a Nan-drop spectrophotometer and a Qubit fluorometer. Libraries were sequenced using 3’ Tag-seq to give 100bp single-end reads at the UT Austin Genomic Sequencing and Analysis Facility (GSAF) on a NovaSeq 6000.

### Data analysis, behavior

Animal coordinates were extracted from raw video files using DeepLabCut v2.1 (DLC, (***Mathis et al., 2018***)). Each video with footage of six individual anemones was cropped into six videos to be analyzed separately. A training dataset was constructed from 3699 images, in which we manually labelled the center-point of individual anemones. The DLC algorithm was trained on this dataset for 600,000 iterations using default settings and the “resnet_50” neural network. DLC output files were analyzed using R code derived from the DLCAnalyzer Github package (https://github.com/ETHZ-INS/DLCAnalyzer, (***Sturman et al., 2020***)); custom code used for this paper is available at this github repository: https://github.com/caberger1/Sensory-Conflict-in-Nematostella-vectensis). Frames with a likelihood less than 0.90 were removed and interpolated, as were frames where the recorded movement was more than 2 cm per frame. An animal was considered “moving” if its speed was at least 0.03 cm s^−1^ for a period of at least 3s (6 frames); any other movement was considered noise and ignored for downstream analysis. For each animal, distance moved was summed into hourly bins based on real-world clock times and normalized to that animal’s maximum hourly movement. This corrects for differences in overall activity due to e.g. body size (as in (***Hendricks et al., 2012; Oren et al., 2015; Tarrant et al., 2019***)). For each experimental group, we averaged the hourly time points to produce a mean time series, which we analyzed alongside the individual time series. Time series were smoothed with a centered moving average in 4h windows.

### Statistics

Two complementary rhythmicity tests were used. The first was a Lomb-Scargle periodogram (LSP) implemented within the R package ‘lomb’, with periods restricted from 20-28h and significance assessed by permutation (n=2000). The second test, the empirical JTK method, was implemented within the BioDare2 web portal (‘eJTK Classic’; https://biodare2.ed.ac.uk/). This approach fits a series of cosinor waves with 24h periods, thus testing for periods of exactly 24h. An LSP test with periods from 10-14h was used to test for circatidal rhythms. In all cases, p-values were adjusted by Benjamin-Hochberg correction, and we chose a conservative FDR cutoff of 0.001 because we were only interested in time series that could confidently be identified as rhythmic. The ‘mFour-Fit’ method implemented in the BioDare2 web portal was used to calculate periods and phases of time series. This is a curve-fitting procedure that is generally the most accurate of the available methods for short time series (***Zielinski et al., 2014***).

The R package ‘CircaCompare’ v0.1.1 was used to test for differences in amplitude and phase of mean time series. CircaCompare uses a cosinor curve-fitting approach and thus gives different estimates of phase and amplitude than those calculated by MFF; however, CircaCompare allows for formal hypothesis testing. The Rayleigh test for circular uniformity (***Rayleigh, 1880***) was used to test whether phases were distributed non-uniformly (implemented in the R package ‘circular’ v0.4). A bootstrapped version of Watson’s non-parametric test (described in (***Fisher et al., 1993; Pewsey et al., 2013***) and implemented in the R package ‘AS.circular’ v0.0.0.9) was used to test whether phase distributions differed in their mean direction; significance was assessed with 9999 bootstraps. The Kruskal-Wallis test followed by Dunn’s test for multiple comparisons were used to test for differences in non-circular means between groups (implemented in R packages ‘stats’ v4.0.3 and ‘dunn.test’ v1.3.5). Levene’s test was used to test for differences in variance (‘car’ v3.0), and paired Wilcoxon rank tests (‘stats’ v4.0.3) were used to test for differences in distance to expected light and temperature phases. Unless otherwise noted, p-values were corrected for multiple testing using the Benjamini-Hochberg procedure.

Linear mixed effects models were used to test for differences in (un-normalized) activity levels between groups, considering two measures of activity: percent time active, and total distance covered. Each hour of activity for each individual was considered a separate observation. We accounted for baseline differences in activity between individuals using random intercepts, and for autocorrelation of residuals using a correlation structure of AR(1); using more complex correlation structures did not affect the results. Mixed models were implemented in the R package ‘nlme’ v3.1, and post-hoc testing was done using ‘emmeans’ v1.6.2.

Time series clustering analysis was done by decomposing smoothed mean time series using wavelet transformation in the R package ‘biwavelet’ v0.20.21, with the smallest scale (period) set to 2 and the largest scale to 36. The ‘wclust’ command was used to construct a distance matrix of the 14 wavelet series, and principal component analysis (PCA) was performed on the distance matrix with ‘FactoMineR’ v2.4. Hierarchical clustering was performed in principal component space with the ‘HCPC’ command using complete linkage. The AU test was used to quantify uncertainty in clustering in the pvclust v2.2 R package (***Suzuki and Shimodaira, 2006***) with 1000 bootstraps, and clustering and PCA analyses were visualized with the ‘factoextra’ package v1.0.7.

### Data analysis, gene expression

Raw reads were trimmed using Cutadapt v3.3 (***Martin, 2011***) with a minimum length of 25 and quality cutoff of 5, and mapped to the Simrbase Nvec200_v1 genome (***Zimmermann et al., 2020***) using Salmon v1.5.1 with “selective alignment (full)” (SAF) (***Srivastava et al., 2020***), a k-mer size of 21, and no length correction (as Tag-seq reads do not have length bias). Tximport v1.16.1 (***Soneson et al., 2016***) was used to import counts into R; counts were then TMM-normalized (***Robinson et al., 2010***) and expressed as counts per million on a log2 scale. Raw reads from (***Oren et al., 2015***) were re-analyzed in the same way as above, except they were length-corrected.

Differential expression (DE) analysis was performed using limma-voom 3.44.3 (***Phipson et al., 2016***) with quality weights (***Liu et al., 2015***), using the following model: “Condition * Light + Temperature + hour”, allowing for linear effects of temperature and total time in the experiment. Low counts were filtered using the command ‘filterByExpr’ in EdgeR v3.30.3 grouped by time point and condition, which retained 12,294/15,290 genes. Rhythmic gene expression was tested using RAIN v1.24.0 ((***Thaben and Westermark, 2014***)), ECHO v4.0.1 (***De los Santos et al., 2020***)), and DryR v1.0.0 ((***Weger et al., 2021***)). In RAIN, search was performed for waveforms with 24 h periods and asymmetries in 4h increments. P-values were corrected using the Bonferroni procedure. Phases and amplitudes of rhythmic genes were calculated using CircaCompare. Rao’s two-sample test for homogeneity (***Jammalamadaka et al., 2021***) was used to test whether groups of phases were drawn from the same distribution (R package ‘TwoCircular’ v1.0), and significance was assessed with 10,000 Monte-Carlo replications. Discriminant analysis of principal components was conducted with the R package ‘adegenet’ v2.1.4 and code modified from (***Dixon et al., 2015***).

Gene ontology enrichment analysis was performed with the GO_MWU R package with default settings (https://github.com/z0on/GO_MWU, (***Wright et al., 2015***)). GO annotations were downloaded from the SimrBase website (https://simrbase.stowers.org/analysis/294, NVEC200.20200813.gff). KEGG enrichment was performed with ‘clusterProfiler’ v3.0.4. To annotate KEGG pathways, protein sequences from an earlier *Nematostella* genome assembly (Nvec1, JGI) were downloaded from the KEGG website (https://www.genome.jp/kegg/ and queried against the SimrBase transcriptome using TBLASTN. SimrBase genes were annotated with the KEGG term of their best-scoring hit in each pathway, using an e-value cutoff of 1 × 10^−10^ and requiring >75% sequence identity. Sliding window enrichment analysis was performed to identify GO and KEGG terms enriched at certain times of day in each treatment. In each 4h sliding window (e.g. ZT0-4, ZT3-6), P-values were calculated (using GO_MWU for GO, and clusterProfiler for KEGG) by comparing genes with phase in that window against all other genes. The enrichment score of a gene at each hourly bin was calculated by averaging the -log10 adjusted p-values from the 3 sliding windows that overlapped the time point (e.g. the score at ZT3 averaged the sliding windows ZT0-4, ZT1-5, and ZT2-6), and a gene was considered enriched if the score at any time point was greater than -log10(0.05).

Weighted Gene Co-expression Network Analysis v1.70 (WGCNA, (***Langfelder and Horvath, 2007***)) was used to cluster rhythmic genes. For the full network constructed from all genes rhythmic in either time series, a soft-thresholding power of 5 was used, and a power of 8 was used for both the Align- and SC-specific networks. In all cases, a signed hybrid adjacency matrix, a minimum module size of 20, biweight midcorrelation (“bicor”), maxPOutliers=0.2, and mergeCutHeight=0.25 were used. Eigengene rhythmicity was assessed using LSPs with 2000 permutations (periods 20-28h). A linear model was used to correlate eigengene expression with light, temperature, and time. Module preservation statistics were calculated within the WGCNA package, and a Sankey plot was constructed using the package ‘networkD3’ v0.4. Enriched GO terms were calculated for each module using GO_MWU with module membership (‘kME’) as input and all other genes as background. KEGG enrichment was performed using a hypergeometric test within ‘clusterProfiler’, as above.

Putative promoter sequences 1kb upstream of transcription start sites were searched for enriched motifs using the HOMER motif discovery tool (findMotifs.pl), searching for motifs of lengths 6, 8, 10, and 12, retaining the top 15 motifs, and using all other promoters as background. Motifs were considered enriched at a p-value of 1 × 10^−5^ for “known” motifs, or 1 × 10^−10^ for *de novo* motifs.

## Acknowledgments

We are particularly grateful to Dr. Carolyn Tepolt for generously providing the aquarium heater, cooler, and temperature controllers, and Dr. Kirstin Meyer-Kaiser for use of her incubator. We also thank Drs. Gregory Fournier, Casey Dunn, Carolyn Tepolt, and Yisrael Schnytzer for helpful discussions.

## Data availability

Raw RNA-seq data have been uploaded to the NCBI Sequence Read Archive (SRA), Bioproject PRJNA826898. R code used for analysis is available at

https://github.com/caberger1/Sensory-Conflict-in-Nematostella-vectensis.

## Appendix 1

### Comparison of RAIN, ECHO, and DryR

90% of rhythmic genes identified by RAIN (p<0.01) had an adjusted ECHO p-value less than 0.05, and 94% of genes with an ECHO p-value less than 0.01 had a RAIN p-value less than 0.05. Log p-values of ECHO and RAIN were highly correlated (r = 0.85). DryR does not calculate p-values, but instead uses model selection to cluster genes based on their expression profile. Briefly, 81% of genes assigned rhythmicity in either time series by DryR were rhythmic in that time series based on RAIN (p<0.01), and 93% had a RAIN p-value less than 0.05; 73% of rhythmic RAIN genes (p<0.01) were assigned to the corresponding cluster(s) in DryR. Thus RAIN and DryR also showed good agreement, although RAIN tended to identify additional rhythmic genes not clustered by DryR. The output of all three programs can be found in ***Figure 5–source data 2***).

## Appendix 2

### Comparison with Oren et al. (2015)

We compared our results to a previous studies of diel/circadian gene expression in Nematostella, ***Oren et al. (2015***) (https://doi.org/10.1038/srep11418), by re-analyzing their raw data using the same software and methods as in the current study, including mapping to the SimrBase genome. ***Oren et al. (2015***) sampled anemones for RNA-seq every 4h for 48h over a LD cycle at a constant temperature 23 °C. This study provides a reference for genes exhibiting rhythmic expression under a light-dark cycle at constant temperature. They used a Fourier analysis to identify 180 genes with rhythmic diel expression, which included putative clock genes *Clock, Helt, CIPC, Cry1a*, and*Cry2* (***Oren et al., 2015***). Using RAIN on their re-analyzed data, we identified 1498 rhythmic genes at p<0.05 (560 genes with p<0.01); we used a less stringent p-value cutoff here because their sample size was much smaller than the sample size in our current study. Despite the differences in genome assembly and statistical methodology between the original paper and our re-analysis, we recapitulated some of the most prominent rhythmic genes in their dataset, including the principal genes hypothesized to play a role in the cnidarian clock (Clock, Cry1a, Cry2, Helt-like, and CIPC). The phases of clock genes in these two experiments are shown in 3.

This re-analysis of this previously published LD experiment demonstrates that core clock gene expression is comparable between an LD cycle at constant temperature, and our study of an LD cycle with an aligned temperature cycle. The phases of putative clock genes (Clock, Helt, CIPC, Cry1a, and Cry2) were approximately the same in the aligned time series and in Oren et al. (within 4h), as were the phases of 2 PAR-bZIP genes. The only exception was Cry1b, which was rhythmic in the current study and not in the re-analyzed dataset (p=0.2).

However, the overlap of all rhythmic genes between these studies was small. Only 477 / 1498 (32%) genes rhythmic in ***Oren et al. (2015***) (p<0.05) had a rhythmic p-value less than 0.05 in the current study (345 had p<0.01). Of these 477 shared genes, just 224 (47%) had phases within 4h of each other, and 108 (23%) shifted by at least 8h. However, there was no overall difference between the phase distributions (Rao’s test, p=0.5). Functional enrichment of the ***Oren et al. (2015***) dataset revealed few similarities with the current study—only a single GO/KEGG term, “Spliceosome”, was expressed at the same time (ZT12-18; ***Figure 5– source data 2***). It is important to note that ***Oren et al. (2015***) used animals derived from a different population than the current study (MD versus MA), and differences in other factors, such as diet and time of feeding, could also affect the circadian transcriptome. Finally, as noted above, the ***Oren et al. (2015***) dataset has a much smaller sample size and thus less power to detect true rhythmic transcripts.

**Figure 3–Figure supplement 1.**
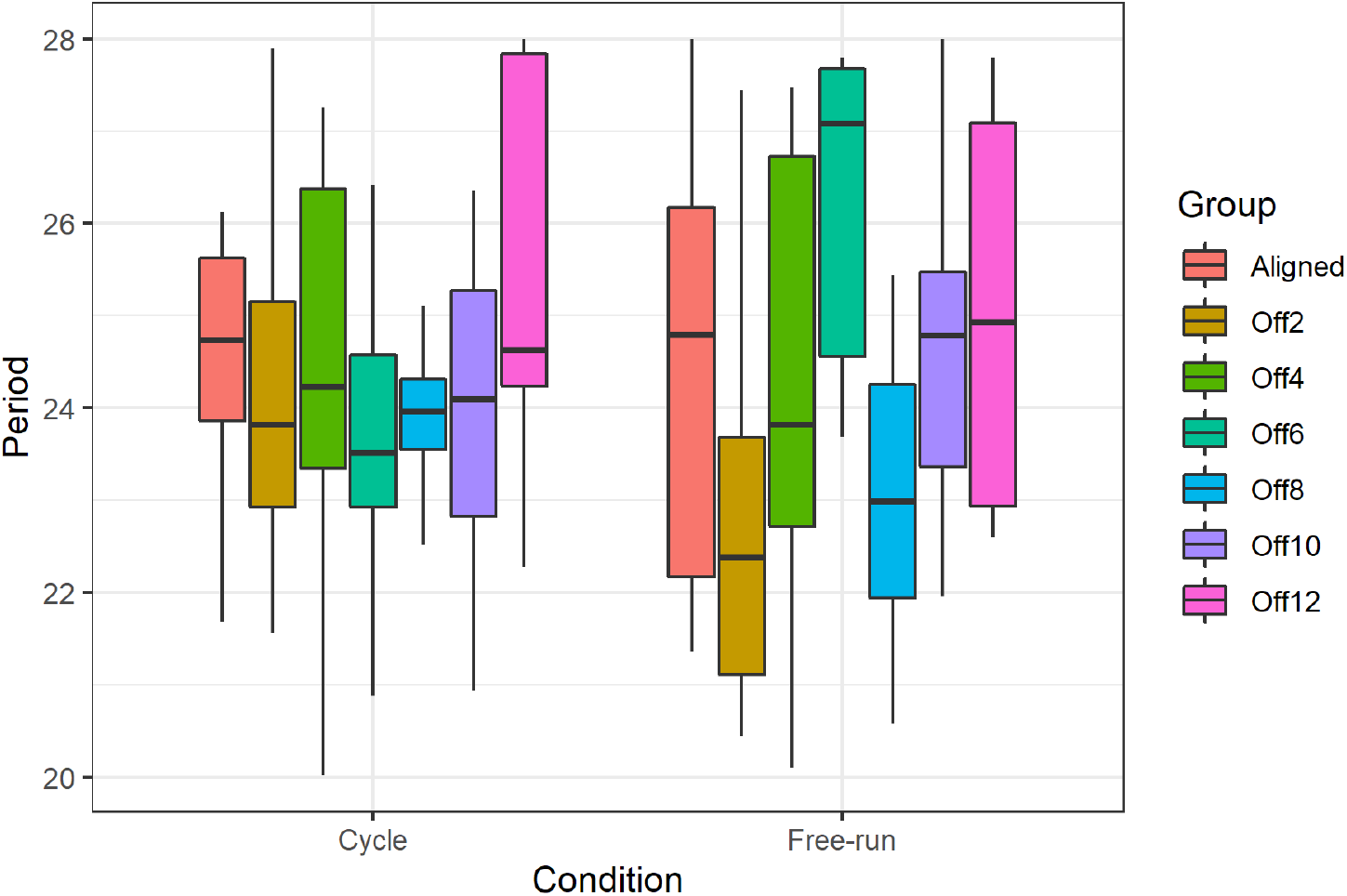
Period lengths of rhythmic individuals. Periods had a wider variance in free-running groups compared to cycling (Levene’s test for variance homogeneity, p=1 × 10^−4^). However, there was no significant difference in mean period between groups (Kruskal-Wallis rank sum test, p=0.096).

**Figure 3–Figure supplement 2.**
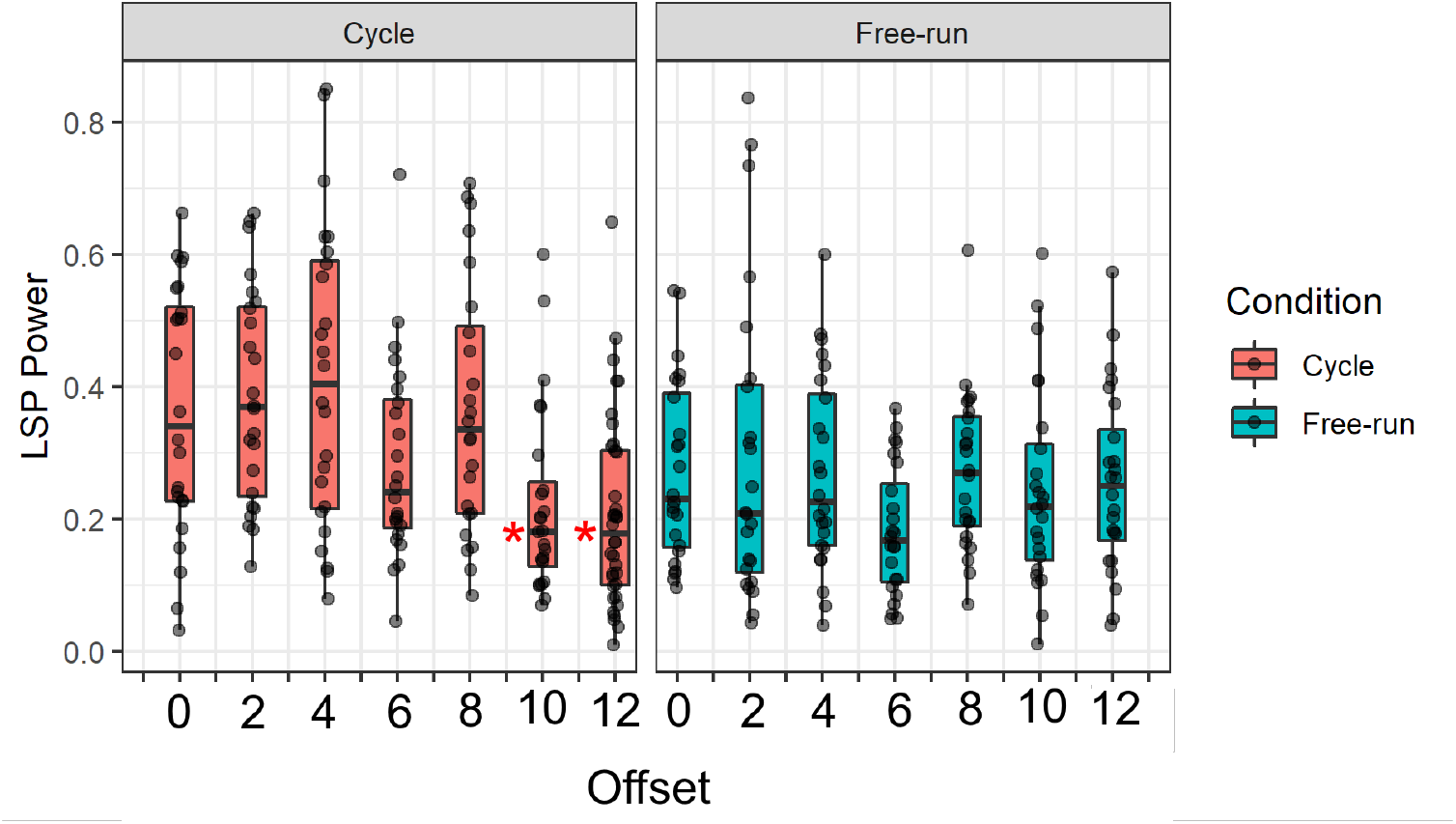
Lomb-Scargle Periodogram (LSP) power across groups. *: Significantly lower than Aligned group (Dunn test, p<0.05).

**Figure 5–Figure supplement 1.**
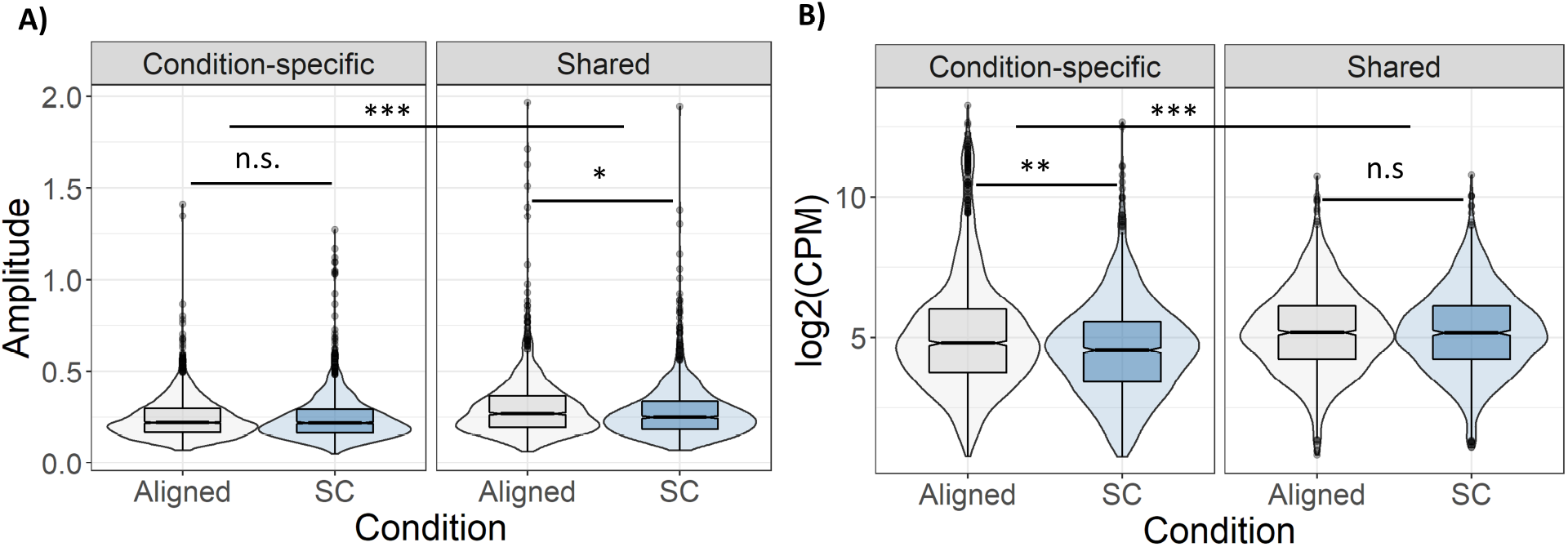
Amplitude and mean expression of rhythmic genes. **A)** Relative amplitude (midpoint to peak) of shared rhythmic genes was reduced during SC. **B)** Mean expression level of condition-specific rhythmic genes was lower during SC. Significance calculated with Wilcoxon signed rank test. n.s.: not significant; *: p<0.01; **: p<1e-5; ***: p<1e-10.

**Figure 6–Figure supplement 1.**
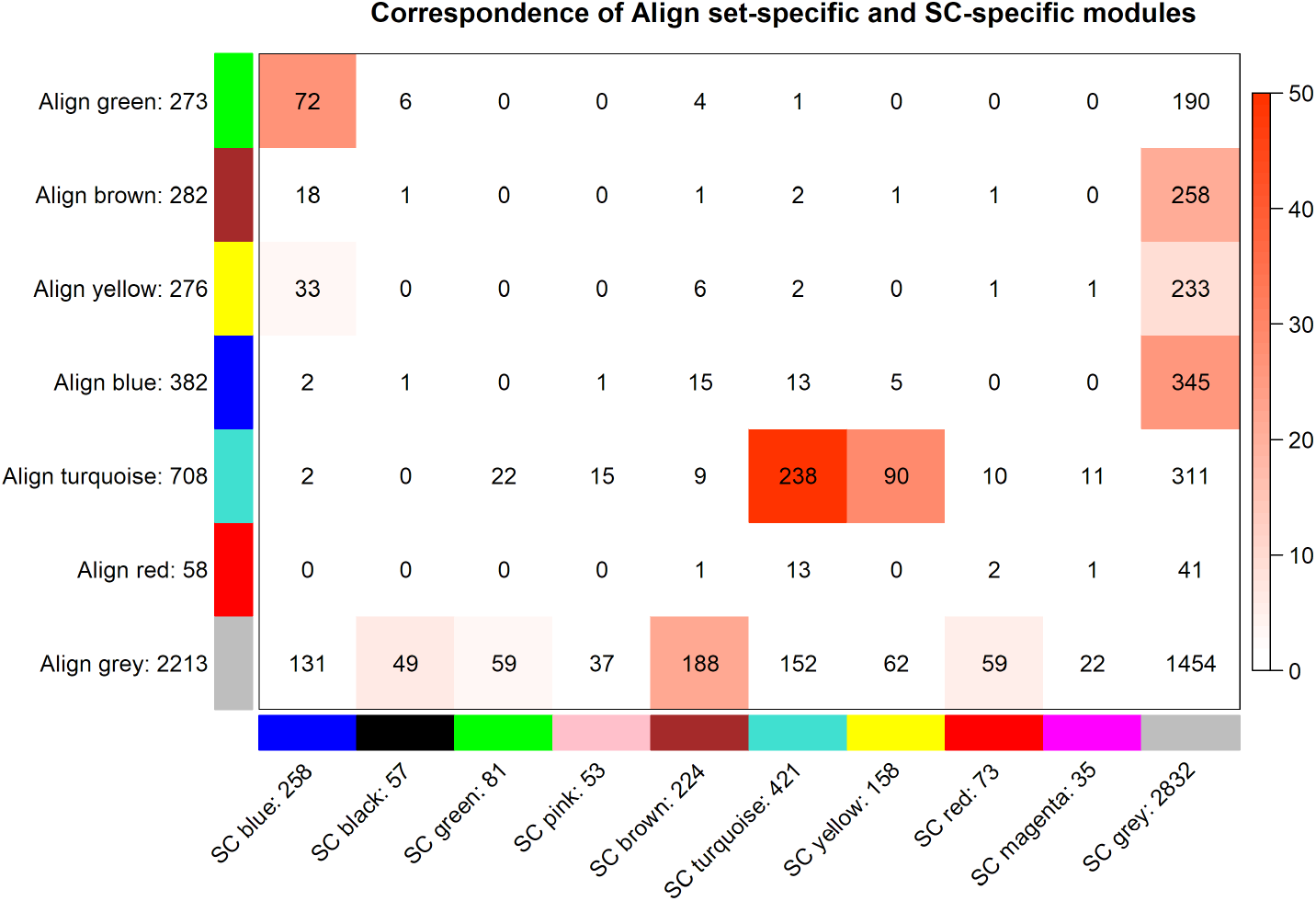
Across-tabulation of Aligned and SC network modules. Numbers along the X and Y axes indicate number of genes in that module; numbers in cells indicate shared genes. Color bar indicates -log10(p-value) of overlap from hypergeometric test.

**Figure 6–Figure supplement 2.**
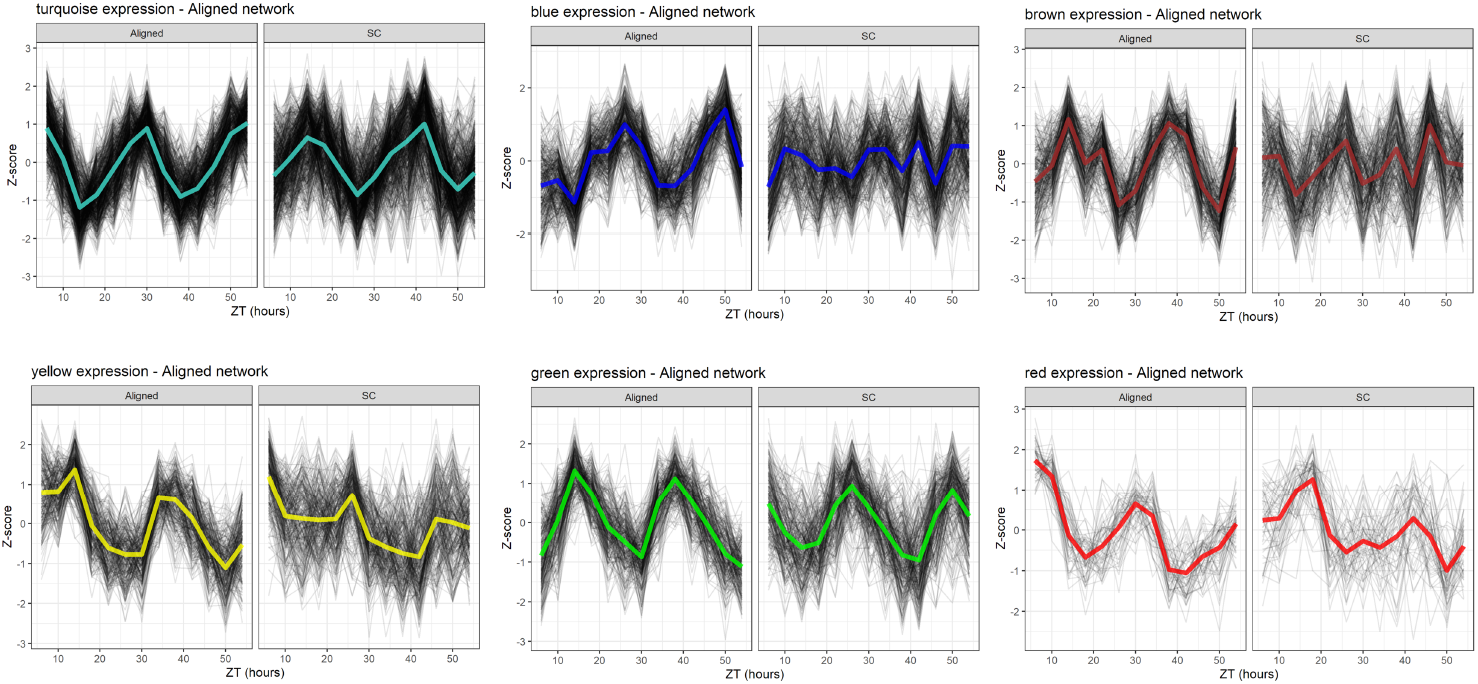
WGCNA module expression, Aligned network. Black lines are Z-scores of the expression of each gene, and colored lines are means of those Z-scores. Left panels show module expression in Aligned samples (where the network was inferred), and right panels show expression in SC samples.

**Figure 6–Figure supplement 3.**
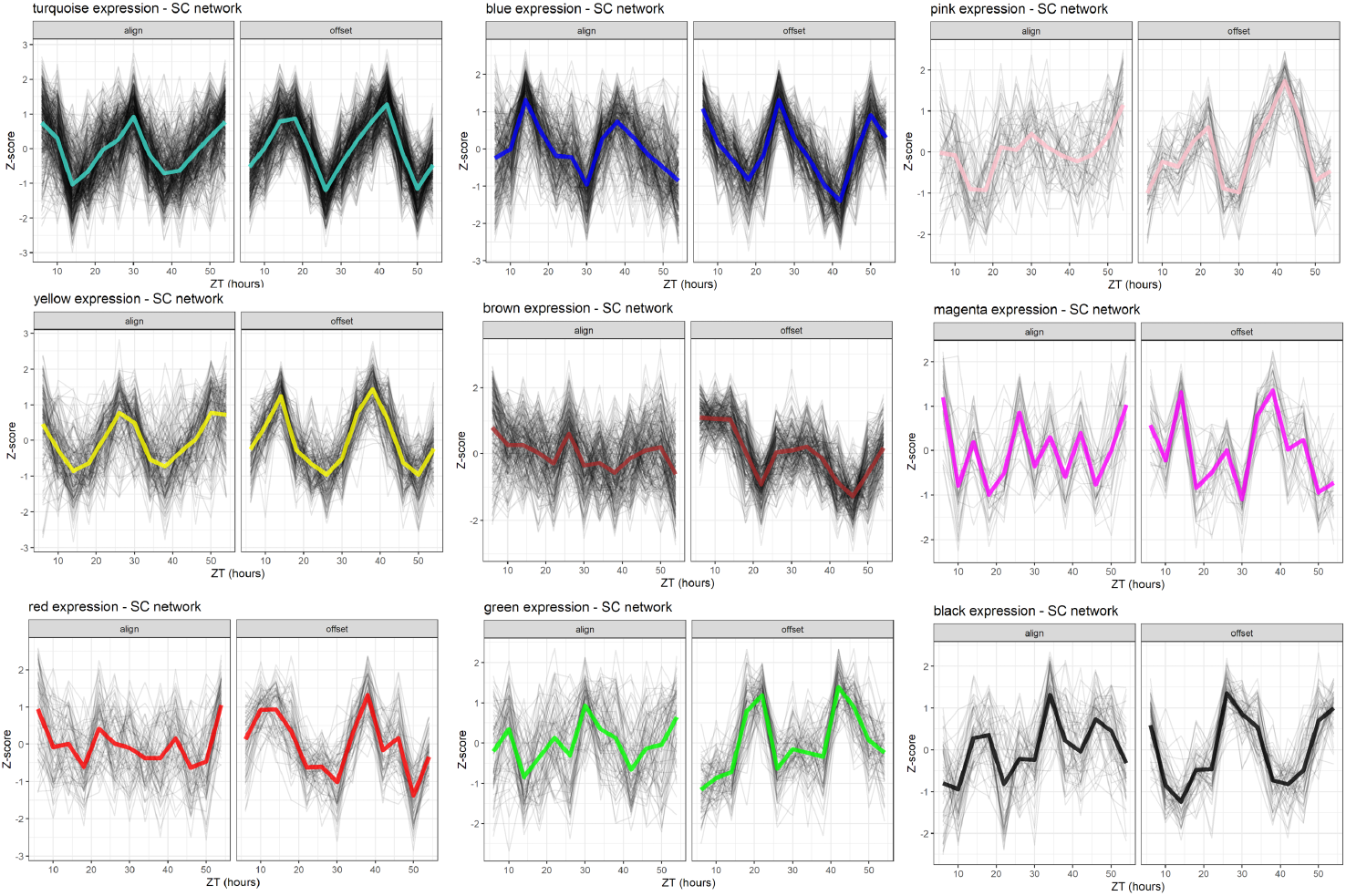
WGCNA module expression, SC network. Black lines are Z-scores of the expression of each gene, and colored lines are means of those Z-scores. Left panels show module expression in Aligned samples, and right panels show expression in SC samples (where the network was inferred).

**Figure 6–Figure supplement 4.**
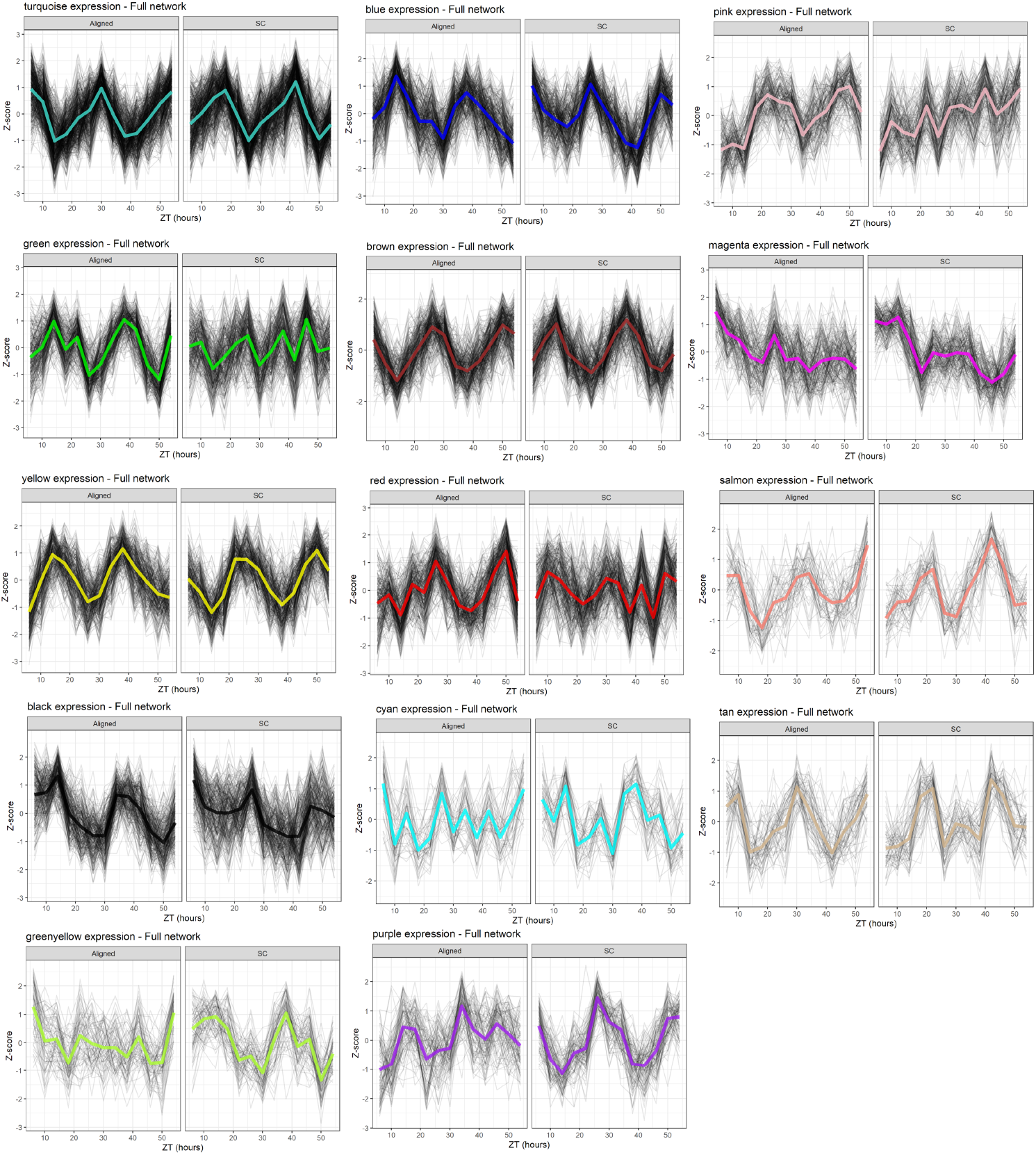
WGCNA module expression, Full network. Black lines are Z-scores of the expression of each gene, and colored lines are means of those Z-scores. Left panels show module expression in Aligned samples, and right panels show expression in SC samples (network was inferred from all samples).

## References

Aschoff J. Exogenous and Endogenous Components in Circadian Rhythms. Cold Spring Harbor Symposia on Quantitative Biology. 1960 Jan; 25:11–28. doi: 10.1101/SQB.1960.025.01.004.

Boothroyd CE, Wijnen H, Naef F, Saez L, Young MW. Integration of Light and Temperature in the Regulation of Circadian Gene Expression in Drosophila. PLoS Genetics. 2007 Apr; 3(4):e54. doi: 10.1371/journal.pgen.0030054.

Bruce VG. Environmental Entrainment of Circadian Rhythms. Cold Spring Harbor Symposia on Quantitative Biology. 1960 Jan; 25:29–48. doi: 10.1101/SQB.1960.025.01.005.

Challet E, Solberg LC, Turek FW. Entrainment in Calorie-Restricted Mice: Conflicting Zeitgebers and Free-Running Conditions. American Journal of Physiology-Regulatory, Integrative and Comparative Physiology. 1998 Jun; 274(6):R1751–R1761. doi: 10.1152/ajpregu.1998.274.6.R1751.

Cheng AH, Bouchard-Cannon P, Hegazi S, Lowden C, Fung SW, Chiang CK, Ness RW, Cheng HYM. SOX2-dependent Transcription in Clock Neurons Promotes the Robustness of the Central Circadian Pacemaker. Cell Reports. 2019 Mar; 26(12):3191–3202.e8. doi: 10.1016/j.celrep.2019.02.068.

Chun LE, Woodruff ER, Morton S, Hinds LR, Spencer RL. Variations in Phase and Amplitude of Rhythmic Clock Gene Expression across Prefrontal Cortex, Hippocampus, Amygdala, and Hypothalamic Paraventricular and Suprachiasmatic Nuclei of Male and Female Rats. Journal of Biological Rhythms. 2015 Oct; 30(5):417–436. doi: 10.1177/0748730415598608.

Currie J, Goda T, Wijnen H. Selective Entrainment of the Drosophila Circadian Clock to Daily Gradients in Environmental Temperature. BMC Biology. 2009 Aug; 7:49. doi: 10.1186/1741-7007-7-49.

Cyran SA, Buchsbaum AM, Reddy KL, Lin MC, Glossop NRJ, Hardin PE, Young MW, Storti RV, Blau J. Vrille, Pdp1, and dClock Form a Second Feedback Loop in the Drosophila Circadian Clock. Cell. 2003 Feb; 112(3):329–341. doi: 10.1016/s0092-8674(03)00074-6.

De los Santos H, Collins EJ, Mann C, Sagan AW, Jankowski MS, Bennett KP, Hurley JM. ECHO: An Application for Detection and Analysis of Oscillators Identifies Metabolic Regulation on Genome-Wide Circadian Output. Bioinformatics. 2020 Feb; 36(3):773–781. doi: 10.1093/bioinformatics/btz617.

Di Rosa V, Frigato E, López-Olmeda JF, Sánchez-Vázquez FJ, Bertolucci C. The Light Wavelength Affects the Ontogeny of Clock Gene Expression and Activity Rhythms in Zebrafish Larvae. PLOS ONE. 2015 Jul; 10(7):e0132235. doi: 10.1371/journal.pone.0132235.

Dibner C, Schibler U, Albrecht U. The Mammalian Circadian Timing System: Organization and Coordination of Central and Peripheral Clocks. Annual Review of Physiology. 2010; 72(1):517–549. doi: 10.1146/annurev-physiol-021909-135821.

Dixon GB, Davies SW, Aglyamova GA, Meyer E, Bay LK, Matz MV. CORAL REEFS. Genomic Determinants of Coral Heat Tolerance across Latitudes. Science (New York, NY). 2015 Jun; 348(6242):1460–1462. doi: 10.1126/science.1261224.

Dubowy C, Sehgal A. Circadian Rhythms and Sleep in Drosophila Melanogaster. Genetics. 2017 Apr; 205(4):1373–1397. doi: 10.1534/genetics.115.185157.

Ehlers CL, Frank E, Kupfer DJ. Social Zeitgebers and Biological Rhythms: A Unified Approach to Understanding the Etiology of Depression. Archives of General Psychiatry. 1988 Oct; 45(10):948–952. doi: 10.1001/archpsyc.1988.01800340076012.

Firth BT, Belan I, Kennaway DJ, Moyer RW. Thermocyclic Entrainment of Lizard Blood Plasma Melatonin Rhythms in Constant and Cyclic Photic Environments. American Journal of Physiology-Regulatory, Integrative and Comparative Physiology. 1999 Dec; 277(6):R1620–R1626. doi: 10.1152/ajpregu.1999.277.6.R1620.

Firth BT, Kennaway DJ. Thermoperiod and Photoperiod Interact to Affect the Phase of the Plasma Melatonin Rhythm in the Lizard, Tiliqua Rugosa. Neuroscience Letters. 1989 Nov; 106(1):125–130. doi: 10.1016/0304-3940(89)90213-9.

Fisher NI, Lewis T, Embleton BJJ. Statistical Analysis of Spherical Data. Cambridge University Press; 1993.

Fowler SG, Cook D, Thomashow MF. Low Temperature Induction of Arabidopsis CBF1, 2, and 3 Is Gated by the Circadian Clock. Plant Physiology. 2005 Mar; 137(3):961–968. doi: 10.1104/pp.104.058354.

Gachon F. Physiological Function of PARbZip Circadian Clock-controlled Transcription Factors. Annals of Medicine. 2007 Jan; 39(8):562–571. doi: 10.1080/07853890701491034.

Gaston KJ, Bennie J, Davies TW, Hopkins J. The Ecological Impacts of Nighttime Light Pollution: A Mechanistic Appraisal. Biological Reviews. 2013; 88(4):912–927. doi: 10.1111/brv.12036.

Glaser FT, Stanewsky R. Temperature Synchronization of the Drosophila Circadian Clock. Current Biology. 2005 Aug; 15(15):1352–1363. doi: 10.1016/j.cub.2005.06.056.

Glossop NRJ, Lyons LC, Hardin PE. Interlocked Feedback Loops Within the Drosophila Circadian Oscillator. Science. 1999 Oct; 286(5440):766–768. doi: 10.1126/science.286.5440.766.

Green CB, Takahashi JS, Bass J. The Meter of Metabolism. Cell. 2008 Sep; 134(5):728–742. doi: 10.1016/j.cell.2008.08.022.

Greenwell BJ, Trott AJ, Beytebiere JR, Pao S, Bosley A, Beach E, Finegan P, Hernandez C, Menet JS. Rhythmic Food Intake Drives Rhythmic Gene Expression More Potently than the Hepatic Circadian Clock in Mice. Cell Reports. 2019 Apr; 27(3):649–657.e5. doi: 10.1016/j.celrep.2019.03.064.

Hardin PE, Panda S. Circadian Timekeeping and Output Mechanisms in Animals. Current Opinion in Neurobiology. 2013 Oct; 23(5):724–731. doi: 10.1016/j.conb.2013.02.018.

Harper REF, Dayan P, Albert JT, Stanewsky R. Sensory Conflict Disrupts Activity of the Drosophila Circadian Network. Cell Reports. 2016 Nov; 17(7):1711–1718. doi: 10.1016/j.celrep.2016.10.029.

Harper REF, Ogueta M, Dayan P, Stanewsky R, Albert JT. D. Journal of Biological Rhythms. 2017 Oct; 32(5):423–432. doi: 10.1177/0748730417724250.

Hart DW, van Jaarsveld B, Lasch KG, Grenfell KL, Oosthuizen MK, Bennett NC. Ambient Temperature as a Strong Zeitgeber of Circadian Rhythms in Response to Temperature Sensitivity and Poor Heat Dissipation Abilities in Subterranean African Mole-Rats. Journal of Biological Rhythms. 2021 Oct; 36(5):461–469. doi: 10.1177/07487304211034287.

Heckmann LH, Sørensen PB, Krogh PH, Sørensen JG. NORMA-Gene: A Simple and Robust Method for qPCR Normalization Based on Target Gene Data. BMC bioinformatics. 2011 Jun; 12:250. doi: 10.1186/1471-2105-12-250.

Hendricks WD, Byrum CA, Meyer-Bernstein EL. Characterization of Circadian Behavior in the Starlet Sea Anemone, Nematostella Vectensis. PLOS ONE. 2012 Oct; 7(10):e46843. doi: 10.1371/journal.pone.0046843.

Hurley JM, Jankowski MS, De los Santos H, Crowell AM, Fordyce SB, Zucker JD, Kumar N, Purvine SO, Robinson EW, Shukla A, Zink E, Cannon WR, Baker SE, Loros JJ, Dunlap JC. Circadian Proteomic Analysis Uncovers Mechanisms of Post-Transcriptional Regulation in Metabolic Pathways. Cell Systems. 2018 Dec; 7(6):613–626.e5. doi: 10.1016/j.cels.2018.10.014.

Jammalamadaka SR, Guerrier S, Mangalam V. A Two-Sample Nonparametric Test for Circular Data–Its Exact Distribution and Performance. Sankhya B. 2021 May; 83(1):140–166. doi: 10.1007/s13571-020-00244-9.

Javier Sánchez-Vázquez F, Zamora S, Madrid JA. Light-Dark and Food Restriction Cycles in Sea Bass: Effect of Conflicting Zeitgebers on Demand-Feeding Rhythms. Physiology & Behavior. 1995 Oct; 58(4):705–714. doi: 10.1016/0031-9384(95)00116-Z.

Kaniewska MM, Vaněčková H, Doležel D, Kotwica-Rolinska J. Light and Temperature Synchronizes Locomotor Activity in the Linden Bug, Pyrrhocoris Apterus. Frontiers in Physiology. 2020 Apr; 11. doi: 10.3389/fphys.2020.00242.

Lague M, Reebs SG. Phase-Shifting the Light–Dark Cycle Influences Food-Anticipatory Activity in Golden Shiners. Physiology & Behavior. 2000 Jul; 70(1):55–59. doi: 10.1016/S0031-9384(00)00246-8.

Lahiri K, Vallone D, Gondi SB, Santoriello C, Dickmeis T, Foulkes NS. Temperature Regulates Transcription in the Zebrafish Circadian Clock. PLoS Biology. 2005 Nov; 3(11). doi: 10.1371/journal.pbio.0030351.

Langfelder P, Horvath S. Eigengene Networks for Studying the Relationships between Co-Expression Modules. BMC Systems Biology. 2007 Nov; 1:54. doi: 10.1186/1752-0509-1-54.

Langfelder P, Luo R, Oldham MC, Horvath S. Is My Network Module Preserved and Reproducible? PLOS Computational Biology. 2011 Jan; 7(1):e1001057. doi: 10.1371/journal.pcbi.1001057.

Leach WB, Reitzel AM. Decoupling Behavioral and Transcriptional Responses to Color in an Eyeless Cnidarian. BMC Genomics. 2020 May; 21(1):361. doi: 10.1186/s12864-020-6766-y.

Lin n, Chou n, Huang n. Priority of Light/Dark Entrainment over Temperature in Setting the Circadian Rhythms of the Prokaryote Synechococcus RF-1. Planta. 1999 Aug; 209(2):202–206. doi: 10.1007/s004250050623.

Liu R, Holik AZ, Su S, Jansz N, Chen K, Leong HS, Blewitt ME, Asselin-Labat ML, Smyth GK, Ritchie ME. Why Weight? Modelling Sample and Observational Level Variability Improves Power in RNA-seq Analyses. Nucleic Acids Research. 2015 Sep; 43(15):e97. doi: 10.1093/nar/gkv412.

Lück S, Thurley K, Thaben PF, Westermark PO. Rhythmic Degradation Explains and Unifies Circadian Transcriptome and Proteome Data. Cell Reports. 2014 Oct; 9(2):741–751. doi: 10.1016/j.celrep.2014.09.021.

Ma D, Li S, Molusky MM, Lin JD. Circadian Autophagy Rhythm: A Link between Clock and Metabolism? Trends in Endocrinology and Metabolism. 2012 Jul; 23(7):319–325. doi: 10.1016/j.tem.2012.03.004.

Mack KL, Jaggard JB, Persons JL, Roback EY, Passow CN, Stanhope BA, Ferrufino E, Tsuchiya D, Smith SE, Slaughter BD, Kowalko J, Rohner N, Keene AC, McGaugh SE. Repeated Evolution of Circadian Clock Dysregulation in Cavefish Populations. PLoS Genetics. 2021 Jul; 17(7):e1009642. doi: 10.1371/journal.pgen.1009642.

Martin M. Cutadapt Removes Adapter Sequences from High-Throughput Sequencing Reads. EMBnetjournal. 2011 May; 17(1):10–12. doi: 10.14806/ej.17.1.200.

Mathis A, Mamidanna P, Cury KM, Abe T, Murthy VN, Mathis MW, Bethge M. DeepLabCut: Markerless Pose Estimation of User-Defined Body Parts with Deep Learning. Nature Neuroscience. 2018 Sep; 21(9):1281–1289. doi: 10.1038/s41593-018-0209-y.

Miyasako Y, Umezaki Y, Tomioka K. Separate Sets of Cerebral Clock Neurons Are Responsible for Light and Temperature Entrainment of Drosophila Circadian Locomotor Rhythms. Journal of Biological Rhythms. 2007 Apr; 22(2):115–126. doi: 10.1177/0748730407299344.

Moyer RW, Firth BT, Lennaway DJ. Effect of Variable Temperatures, Darkness and Light on the Secretion of Melatonin by Pineal Explants in the Gecko, Christinus Marmoratus. Brain Research. 1997 Feb; 747(2):230–235. doi: 10.1016/S0006-8993(96)01266-8.

Nedeltcheva AV, Scheer FAJL. Metabolic Effects of Sleep Disruption, Links to Obesity and Diabetes. Current opinion in endocrinology, diabetes, and obesity. 2014 Aug; 21(4):293–298. doi: 10.1097/MED.0000000000000082.

Nikhil KL, Goirik G, Ratna K, Sharma VK. Role of Temperature in Mediating Morning and Evening Emergence Chronotypes in Fruit Flies Drosophila Melanogaster. Journal of Biological Rhythms. 2014 Dec; 29(6):427–441. doi: 10.1177/0748730414553797.

Oren M, Tarrant AM, Alon S, Simon-Blecher N, Elbaz I, Appelbaum L, Levy O. Profiling Molecular and Behavioral Circadian Rhythms in the Non-Symbiotic Sea Anemone Nematostella Vectensis. Scientific Reports. 2015 Jun; 5:11418. doi: 10.1038/srep11418.

Pácha J, Balounová K, Soták M. Circadian Regulation of Transporter Expression and Implications for Drug Disposition. Expert Opinion on Drug Metabolism & Toxicology. 2021 Apr; 17(4):425–439. doi: 10.1080/17425255.2021.1868438.

Partch CL, Green CB, Takahashi JS. Molecular Architecture of the Mammalian Circadian Clock. Trends in Cell Biology. 2014 Feb; 24(2):90–99. doi: 10.1016/j.tcb.2013.07.002.

Pewsey A, Neuhäuser M, Ruxton GD. Circular Statistics in R. OUP Oxford; 2013.

Phipson B, Lee S, Majewski IJ, Alexander WS, Smyth GK. ROBUST HYPERPARAMETER ESTIMATION PROTECTS AGAINST HYPERVARIABLE GENES AND IMPROVES POWER TO DETECT DIFFERENTIAL EXPRESSION. The annals of applied statistics. 2016 Jun; 10(2):946–963. doi: 10.1214/16-AOAS920.

Rayleigh L. XII. On the Resultant of a Large Number of Vibrations of the Same Pitch and of Arbitrary Phase. The London, Edinburgh, and Dublin Philosophical Magazine and Journal of Science. 1880 Aug; 10(60):73–78. doi: 10.1080/14786448008626893.

Reierth E, Stokkan KA. Dual Entrainment by Light and Food in the Svalbard Ptarmigan (Lagopus Mutus Hyperboreus). Journal of Biological Rhythms. 1998 Oct; 13(5):393–402. doi: 10.1177/074873049801300504.

Reitzel AM, Behrendt L, Tarrant AM. Light Entrained Rhythmic Gene Expression in the Sea Anemone Nematostella Vectensis: The Evolution of the Animal Circadian Clock. PLOS ONE. 2010 Sep; 5(9):e12805. doi: 10.1371/journal.pone.0012805.

Reitzel AM, Tarrant AM. Nuclear Receptor Complement of the Cnidarian Nematostella Vectensis: Phylogenetic Relationships and Developmental Expression Patterns. BMC Evolutionary Biology. 2009 Sep; 9(1):230. doi: 10.1186/1471-2148-9-230.

Reitzel AM, Tarrant AM, Levy O. Circadian Clocks in the Cnidaria: Environmental Entrainment, Molecular Regulation, and Organismal Outputs. Integrative and Comparative Biology. 2013 Jul; 53(1):118–130. doi: 10.1093/icb/ict024.

Rey G, Valekunja UK, Feeney KA, Wulund L, Milev NB, Stangherlin A, Ansel-Bollepalli L, Velagapudi V, O’Neill JS, Reddy AB. The Pentose Phosphate Pathway Regulates the Circadian Clock. Cell Metabolism. 2016 Sep; 24(3):462–473. doi: 10.1016/j.cmet.2016.07.024.

Rivas GBS, Bauzer LGSdR, Meireles-Filho ACA. “The Environment Is Everything That Isn’t Me”: Molecular Mechanisms and Evolutionary Dynamics of Insect Clocks in Variable Surroundings. Frontiers in Physiology. 2016; 6. doi: 10.3389/fphys.2015.00400.

Rivas GBS, Teles-de-Freitas R, Pavan MG, Lima JBP, Peixoto AA, Bruno RV. Effects of Light and Temperature on Daily Activity and Clock Gene Expression in Two Mosquito Disease Vectors. Journal of Biological Rhythms. 2018 Jun; 33(3):272–288. doi: 10.1177/0748730418772175.

Robinson MD, McCarthy DJ, Smyth GK. edgeR: A Bioconductor Package for Differential Expression Analysis of Digital Gene Expression Data. Bioinformatics. 2010 Jan; 26(1):139–140. doi: 10.1093/bioinformatics/btp616.

Roenneberg T, Merrow M. The Circadian Clock and Human Health. Current biology: CB. 2016 May; 26(10):R432–443. doi: 10.1016/j.cub.2016.04.011.

Ruijter JM, Ramakers C, Hoogaars WMH, Karlen Y, Bakker O, van den Hoff MJB, Moorman AFM. Amplification Effciency: Linking Baseline and Bias in the Analysis of Quantitative PCR Data. Nucleic Acids Research. 2009 Apr; 37(6):e45–e45. doi: 10.1093/nar/gkp045.

Sachkova MY, Macrander J, Surm JM, Aharoni R, Menard-Harvey SS, Klock A, Leach WB, Reitzel AM, Moran Y. Some like It Hot: Population-Specific Adaptations in Venom Production to Abiotic Stressors in a Widely Distributed Cnidarian. BMC Biology. 2020 Sep; 18(1):121. doi: 10.1186/s12915-020-00855-8.

Salter MG, Franklin KA, Whitelam GC. Gating of the Rapid Shade-Avoidance Response by the Circadian Clock in Plants. Nature. 2003 Dec; 426(6967):680–683. doi: 10.1038/nature02174.

Schmal C, Ono D, Myung J, Pett JP, Honma S, Honma KI, Herzel H, Tokuda IT. Weak Coupling between Intracellular Feedback Loops Explains Dissociation of Clock Gene Dynamics. PLOS Computational Biology. 2019 Sep; 15(9):e1007330. doi: 10.1371/journal.pcbi.1007330.

Sebé-Pedrós A, Saudemont B, Chomsky E, Plessier F, Mailhé MP, Renno J, Loe-Mie Y, Lifshitz A, Mukamel Z, Schmutz S, Novault S, Steinmetz PRH, Spitz F, Tanay A, Marlow H. Cnidarian Cell Type Diversity and Regulation Revealed by Whole-Organism Single-Cell RNA-seq. Cell. 2018 May; 173(6):1520–1534.e20. doi: 10.1016/j.cell.2018.05.019.

Simion P, Philippe H, Baurain D, Jager M, Richter DJ, Di Franco A, Roure B, Satoh N, Quéinnec É, Ereskovsky A, Lapébie P, Corre E, Delsuc F, King N, Wörheide G, Manuel M. A Large and Consistent Phylogenomic Dataset Supports Sponges as the Sister Group to All Other Animals. Current Biology. 2017 Apr; 27(7):958–967. doi: 10.1016/j.cub.2017.02.031.

Somers DE, Webb AA, Pearson M, Kay SA. The Short-Period Mutant, Toc1-1, Alters Circadian Clock Regulation of Multiple Outputs throughout Development in Arabidopsis Thaliana. Development (Cambridge, England). 1998 Feb; 125(3):485–494. doi: 10.1242/dev.125.3.485.

Somers J, Harper REF, Albert JT. How Many Clocks, How Many Times? On the Sensory Basis and Computational Challenges of Circadian Systems. Frontiers in Behavioral Neuroscience. 2018; 12. doi: 10.3389/fn-beh.2018.00211.

Soneson C, Love MI, Robinson MD. Differential Analyses for RNA-Seq: Transcript-level Estimates Improve Gene-Level Inferences. F1000Research. 2016 Feb; 4. doi: 10.12688/f1000research.7563.2.

Srivastava A, Malik L, Sarkar H, Zakeri M, Almodaresi F, Soneson C, Love MI, Kingsford C, Patro R. Alignment and Mapping Methodology Influence Transcript Abundance Estimation. Genome Biology. 2020 Sep; 21(1):239. doi: 10.1186/s13059-020-02151-8.

Sturman O, von Ziegler L, Schläppi C, Akyol F, Privitera M, Slominski D, Grimm C, Thieren L, Zerbi V, Grewe B, Bohacek J. Deep Learning-Based Behavioral Analysis Reaches Human Accuracy and Is Capable of Outperforming Commercial Solutions. Neuropsychopharmacology. 2020 Oct; 45(11):1942–1952. doi: 10.1038/s41386-020-0776-y.

Suzuki R, Shimodaira H. Pvclust: An R Package for Assessing the Uncertainty in Hierarchical Clustering. Bioin-formatics. 2006 Jun; 22(12):1540–1542. doi: 10.1093/bioinformatics/btl117.

Tarrant AM, Helm RR, Levy O, Rivera HE. Environmental Entrainment Demonstrates Natural Circadian Rhythmicity in the Cnidarian Nematostella Vectensis. The Journal of Experimental Biology. 2019 Nov; 222(21):jeb205393. doi: 10.1242/jeb.205393.

Thaben PF, Westermark PO. Detecting Rhythms in Time Series with RAIN. Journal of Biological Rhythms. 2014 Dec; 29(6):391–400. doi: 10.1177/0748730414553029.

Thurley K, Herbst C, Wesener F, Koller B, Wallach T, Maier B, Kramer A, Westermark PO. Principles for Cir-cadian Orchestration of Metabolic Pathways. Proceedings of the National Academy of Sciences. 2017 Feb; 114(7):1572–1577. doi: 10.1073/pnas.1613103114.

Turek FW, Joshu C, Kohsaka A, Lin E, Ivanova G, McDearmon E, Laposky A, Losee-Olson S, Easton A, Jensen DR, Eckel RH, Takahashi JS, Bass J. Obesity and Metabolic Syndrome in Circadian Clock Mutant Mice. Science (New York, NY). 2005 May; 308(5724):1043–1045. doi: 10.1126/science.1108750.

Valenciano AI, Alonso-Gómez AL, Alonso-Bedate M, Delgado MJ. Effect of Constant and Fluctuating Temperature on Daily Melatonin Production by Eyecups from Rana Perezi. Journal of Comparative Physiology B, Bio-chemical, Systemic, and Environmental Physiology. 1997 Apr; 167(3):221–228. doi: 10.1007/s003600050068.

Warner JR. The Economics of Ribosome Biosynthesis in Yeast. Trends in Biochemical Sciences. 1999 Nov; 24(11):437–440. doi: 10.1016/s0968-0004(99)01460-7.

Watari Y, Tanaka K. Interacting Effect of Thermoperiod and Photoperiod on the Eclosion Rhythm in the Onion Fly, Delia Antiqua Supports the Two-Oscillator Model. Journal of Insect Physiology. 2010 Sep; 56(9):1192–1197. doi: 10.1016/j.jinsphys.2010.03.022.

Weger BD, Gobet C, David FPA, Atger F, Martin E, Phillips NE, Charpagne A, Weger M, Naef F, Gachon F. Systematic Analysis of Differential Rhythmic Liver Gene Expression Mediated by the Circadian Clock and Feeding Rhythms. Proceedings of the National Academy of Sciences. 2021 Jan; 118(3). doi: 10.1073/pnas.2015803118.

Wright RM, Aglyamova GV, Meyer E, Matz MV. Gene Expression Associated with White Syndromes in a Reef Building Coral, Acropora Hyacinthus. BMC Genomics. 2015 May; 16(1):371. doi: 10.1186/s12864-015-1540-2.

Yeung J, Naef F. Rhythms of the Genome: Circadian Dynamics from Chromatin Topology, Tissue-Specific Gene Expression, to Behavior. Trends in Genetics. 2018 Dec; 34(12):915–926. doi: 10.1016/j.tig.2018.09.005.

Yoshida T, Murayama Y, Ito H, Kageyama H, Kondo T. Nonparametric Entrainment of the in Vitro Circadian Phosphorylation Rhythm of Cyanobacterial KaiC by Temperature Cycle.Proceedings of the National Academy of Sciences of the United States of America. 2009 Feb; 106(5):1648–1653. doi: 10.1073/pnas.0806741106.

Yoshii T, Hermann C, Helfrich-Förster C. Cryptochrome-Positive and -Negative Clock Neurons in Drosophila Entrain Differentially to Light and Temperature. Journal of Biological Rhythms. 2010 Dec; 25(6):387–398. doi: 10.1177/0748730410381962.

Zhang R, Lahens NF, Ballance HI, Hughes ME, Hogenesch JB. A Circadian Gene Expression Atlas in Mammals: Implications for Biology and Medicine. Proceedings of the National Academy of Sciences. 2014 Nov; 111(45):16219–16224. doi: 10.1073/pnas.1408886111.

Zielinski T, Moore AM, Troup E, Halliday KJ, Millar AJ. Strengths and Limitations of Period Estimation Methods for Circadian Data. PLOS ONE. 2014 May; 9(5):e96462. doi: 10.1371/journal.pone.0096462.

Zimmermann B, Robb SMC, Genikhovich G, Fropf WJ, Weilguny L, He S, Chen S, Lovegrove-Walsh J, Hill EM, Ragkousi K, Praher D, Fredman D, Moran Y, Gibson MC, Technau U, Sea Anemone Genomes Reveal Ancestral Metazoan Chromosomal Macrosynteny; 2020. doi: 10.1101/2020.10.30.359448.

